# HERCULES: a deep-learning framework for RNA-binding region localization and mutation-effect prediction

**DOI:** 10.64898/2026.03.17.712455

**Authors:** Jonathan Fiorentino, Michele Monti, Alexandros Armaos, Dimitrios Miltiadis-Vrachnos, Riccardo Delli Ponti, Lorenzo Di Rienzo, Veronica Venturi, Teresa Botta Orfila, Eulalia Marti, Gian Gaetano Tartaglia

**Author notes:** These authors contributed equally to this work (Jonathan Fiorentino, Michele Monti).

## Abstract

RNA-binding proteins regulate essential steps of RNA metabolism, yet identifying RNA-interacting regions and prioritizing sequence variants that alter binding remain challenging from protein sequence alone. Here we present *cat*RAPID *signature* 2.0 HERCULES (**H**ybrid fram**E**wo**R**k for RNA-binding domain lo**C**alization and m**U**tation ana**L**ysis using physicochemical and languag**E** model**S)**, an approach combining a fine-tuned ProteinBERT model with residue-level physicochemical descriptors to predict global RNA-binding propensity, localize interaction sites and estimate the impact of amino acid substitutions. The language model captures domain-scale sequence context, whereas the physicochemical module provides sensitivity to local residue changes. On an independent benchmark, HERCULES discriminates RBPs from non-RBPs with an AUROC of 0.86. At residue resolution, it enriches high-scoring positions within annotated interaction regions across canonical, non-canonical and putative domain classes, while maintaining strong performance on an independent InterPro benchmark. Protein microarray experiments using disease-associated repeat RNAs (CGG, CAG and interrupted CAACAG repeats) independently support multiple non-catalogued candidate RBPs prioritized by HERCULES. Using curated UniProt variants associated with impaired RNA binding, HERCULES also prioritizes deleterious substitutions from predicted changes in binding propensity. By integrating protein-level classification, residue localization, variant-effect prediction and experimental validation, HERCULES provides an interpretable framework for prioritizing RNA-binding proteins, interaction regions and mutation-sensitive sites. HERCULES is freely available at https://tools.tartaglialab.com/hercules and as an open-source Python package at https://github.com/tartaglialabIIT/hercules.git.

## Introduction

RNA-binding proteins (RBPs) regulate essentially all stages of RNA metabolism, including splicing, localization, stability, and translation, and their dysfunction has been implicated in cancer, neurodegeneration, cardiovascular disease, and other pathologies (1–3). As a result, RBPs are increasingly recognized as important biomarkers and therapeutic targets, motivating substantial interest in computational methods that can identify RNA-binding proteins and characterize their interactions directly from sequence (4, 5).

Over the past decade, numerous computational approaches have been developed to identify RBPs. Early methods relied on manually engineered features derived from amino acid composition, physicochemical properties, and evolutionary profiles, typically coupled to classical machine-learning classifiers (6–9). More recently, deep-learning architectures and protein language model (PLM)–based frameworks have substantially improved performance by leveraging large-scale sequence representations and contextual information (10, 11). Among these, HydRA represents a state-of-the-art hybrid approach that integrates a PLM with protein–protein interaction networks and sequence-derived features, achieving strong performance in global RBP classification across benchmark datasets (11). Importantly, HydRA also introduced an occlusion-based strategy to associate global RNA-binding predictions with specific regions along the protein sequence, partially bridging protein-level classification and residue-level interpretation.

Despite substantial progress in identifying RBPs, localizing RNA-binding regions within proteins and assessing how sequence variation may alter RNA-binding propensity remain open challenges. Experimental determination of protein–RNA interfaces relies on structural approaches such as X-ray crystallography, cryo-EM and NMR spectroscopy (12), which can provide residue-level information but are low-throughput and available for only a limited and biased subset of protein–RNA interactions. This has motivated the development of sequence-based predictors of RNA-binding interfaces (13–15). These methods typically aim to identify RNA-interacting residues, often using annotations derived from structurally resolved protein–RNA complexes. More recent approaches have addressed some of these limitations: for example, HybridRNAbind combines predictors trained on structure-annotated and disorder-annotated RNA-binding residues (16), while GPSite uses PLM-derived sequence embeddings together with ESMFold-predicted structures and geometric deep learning to predict binding residues for RNA and other ligands (17). Nevertheless, these methods are primarily optimized for RNA-contact or binding-site prediction rather than for recovering complete RBD-scale RNA-binding propensity profiles. As a result, they may provide sparse or contact-like predictions that do not necessarily coincide with full functional domains, particularly in intrinsically disordered regions or non-canonical RBDs.

On the other hand, HydRA adopts a different strategy by coupling global RNA-binding classification with occlusion-based mapping along the protein sequence, partially mitigating the reliance on structural annotations (11); nevertheless, its dependence on sliding-window perturbations inherently constrains spatial resolution and reduces sensitivity to subtle residue-level physicochemical effects.

Structure-based and predicted-structure-based approaches remain highly valuable, especially when reliable protein–RNA complexes or accurate predicted structures are available. At the same time, independent evaluations indicate that current structure-based approaches still struggle to reliably predict many protein–RNA complexes, particularly those involving intrinsically disordered regions, non-canonical interfaces, or flexible RNAs (18), underscoring the importance of complementary physicochemical models that explicitly capture local biochemical determinants of RNA binding (19). These limitations reflect both the scarcity and bias of experimentally solved protein–RNA structures and the inherent conformational heterogeneity of RNA (20), limiting the applicability of purely structure-driven pipelines for systematic RBD identification or mutation effect prediction.

Here we present *cat*RAPID *signature* 2.0 HERCULES (**H**ybrid fram**E**wo**R**k for RNA-binding domain lo**C**alization and m**U**tation ana**L**ysis using physicochemical and languag**E** model**S)**, a computational framework designed to address this gap. HERCULES integrates two complementary views of protein sequence: first, a protein language model is fine-tuned to discriminate RBPs from non-RBPs, and its global attention layers are leveraged to derive residue-level profiles that highlight regions likely to participate in RNA binding. This component captures global, sequence-wide constraints such as domain architecture and long-range dependencies encoded in PLM representations and high attention residues (21–24); second, an explicit physicochemical module models RNA-binding propensity at residue resolution based on local properties, enabling sensitivity to small numbers of amino acid substitutions and providing a natural mechanism for mutation effect prediction (25).

By combining a PLM-derived domain profile with a mutation-sensitive physicochemical profile, HERCULES enables the simultaneous identification of RNA-binding domains, prediction of global RNA-binding propensity, and mutation-associated propensity changes. This unified framework provides mechanistic insight into protein–RNA interactions while remaining fully sequence-based, making it broadly applicable across species and protein families. In addition to advancing computational RBP annotation, the ability to localize RNA-binding regions and predict mutation effects offers a principled starting point for rational design strategies, including the development and optimization of RNA aptamers targeting specific protein surfaces (1, 3, 26, 27). HERCULES is available both as a user-friendly web server (https://tools.tartaglialab.com/hercules) and a downloadable Python package (https://github.com/tartaglialabIIT/hercules.git).

## Material and Methods

### Datasets

#### Training and test datasets for binary classification

We compiled a list of 3115 human RNA-binding proteins (RBPs) curated for the scRAPID-web resource (28, 29). The list of RBPs was constructed by combining proteins from the RBP2GO database with a score ≥10 (30) and the catRAPID omics v2.0 RBP libraries (31). To reduce redundancy, sequences were clustered using CD-HIT (32) at 50% sequence identity (parameters: -c 0.5 -n 3), resulting in 2675 non-redundant human RBPs. Non-RBPs were obtained combining 1232 experimentally validated negative proteins from HeLa lysate (33), previously employed by catRAPID signature (9), with a random sample of 1443 proteins, yielding a balanced dataset. These additional non-RBPs were randomly sampled from the intersection of three datasets:

- A curated list of human non-RBPs (34);

- The negative protein set from catGRANULE 2.0 ROBOT (25), consisting of proteins not undergoing liquid-liquid phase separation (LLPS), as LLPS proteins are often capable of binding RNAs;

- A list of human non-RBPs from the RBP2GO database (30).

This final dataset was first split into training and independent test sets using an 80/20 ratio. The training partition was subsequently split into training and validation subsets for model selection, early stopping and threshold selection.

### RBP2GO database

RBP2GO is a database of RNA-binding proteins covering 22 species, comprising more than 22000 RBP candidates (30). We used RBP2GO to evaluate the ability of HERCULES to predict, directly from protein sequence, whether a protein is an RBP. Annotated RBPs for nine species were downloaded from the RBP2GO web resource (https://rbp2go.dkfz.de/).

### Pfam RNA-binding region benchmark

We compiled a comprehensive dataset of RNA-binding domains (RBDs) by integrating 67 canonical, 34 non-classical, and 37 putative domains obtained from the Pfam database (35), based on previously established annotations from the catRAPID signature resource (9). RNA Binding Domain occurrences were identified in the reviewed human proteome from UniProt by scanning sequences with HMMER v3.3 (hmmscan) against the Pfam profile HMM library (Pfam 33.1). Matches were accepted using Pfam model-specific GA (gathering) thresholds (*--cut_ga*). This yielded 672 canonical RBD hits, 562 non-classical RBD hits, and 471 putative/unknown RBD hits. Merging RBD annotations across entries corresponding to the same protein resulted in 579 proteins with canonical RBDs, 533 with non-canonical RBDs, and 399 with putative RBDs.To minimize overlap with the HERCULES training set, validation proteins were compared against all training sequences using MMseqs2 easy-search. Proteins were excluded if their best training-set hit had sequence identity ≥30% and bidirectional alignment coverage ≥80%, where coverage was defined as min (query coverage, target coverage). After this filtering, the final non-redundant validation set contained 604 proteins: 215 with canonical RBDs, 248 with non-classical RBDs and 180 with putative RBDs. Because 33 proteins contained domains from more than one class, class-stratified analyses were performed on 643 protein–RBD-class entries: 215 canonical, 248 non-classical and 180 putative.

Canonical RBDs in the dataset include well-characterized domains such as the RNA-recognition motif (RRM), double-stranded RNA-binding domain (dsRBD), K-homology (KH) domain, RGG box, and Pumilio/FBF (PUM) domain. Non-classical and putative RBDs were identified based on experimental studies indicating RNA-binding potential beyond the canonical set (33, 36, 37). Non-classical domains may interact with RNA in less conventional ways, while putative domains are predicted or experimentally suggested to bind RNA but lack extensive characterization.

For global residue-level validation, we combined proteins containing canonical, non-classical, or putative RBDs into a single benchmark set, merging multiple domain occurrences within the same protein into a unified residue-level annotation (per-residue maximum across domain-specific Pfam masks). Residue-level performance was evaluated on the 604-protein non-redundant set described above.

### InterPro RNA-binding region benchmark

We assembled an independent residue-level validation set from non-Pfam InterPro member-database annotations (InterPro release 109.0). Starting from the human Swiss-Prot reviewed proteome (UniProt, 26 February 2026), we extracted region coordinates from InterPro and from UniProt DOMAIN/REGION/REPEAT/MOTIF features with database cross-references. Only annotations supported by non-Pfam member databases (principally PROSITE, SMART, CDD, SUPERFAMILY, Gene3D, and PANTHER) were retained; Pfam-only matches were excluded. Candidate regions were filtered using predefined keyword rules applied to InterPro and UniProt annotation text and linked GO terms. Regions were accepted if they indicated RNA binding or known RNA-binding domain types (for example RRM, KH, dsRBD, DEAD-box helicase, RGG/RG-rich, LSM/Sm, PUF, La motif, cold-shock domain, or snRNP/splicing-related terms) and rejected if they indicated non-specific binding (for example DNA-only, ATP-binding, generic helicase or zinc finger without RNA context, or ribosomal labels without domain-level RNA-binding evidence). Overlapping or adjacent accepted intervals were merged, and residue labels were set to 1 within merged regions and 0 elsewhere.

Proteins with ≥30% sequence identity to the HERCULES training set were removed using the same criterion as the Pfam benchmark. The final strict non-Pfam set comprised 140 proteins, 179 merged regions, and ∼30,100 labelled residues (median ∼39% of each sequence). 76 of these proteins also appear in the Pfam set, but labels were defined independently from non-Pfam sources. Predictors were evaluated on these masks using the same residue-level metrics as the Pfam benchmark.

### Protein–RNA complexes from the Protein Data Bank

We retrieved experimentally resolved 3D structures of protein–RNA complexes from the Protein Data Bank (PDB). The initial structural dataset of protein–RNA complexes was originally extracted from the data described by Bellucci et al (38) and Muppirala et al (39). Starting from 369 non-ribosomal complexes, we defined interacting protein residues and RNA nucleotides using a full-atom distance cutoff of 4 Å. Hence, complexes with fewer than 5 interacting protein residues or fewer than 3 interacting nucleotides were excluded, in order to focus on interfaces with an adequate level of complexity. In addition, to avoid working with small polypeptidic chains (or not entirely solved proteins), only proteins with less than 50% of their residues involved in RNA interaction were retained. After applying these filtering criteria, the structural dataset comprised 266 protein–RNA complexes. Finally, we merged entries containing the same protein and different RNA binding residues annotated, obtaining 176 proteins.

### Mutations affecting protein RNA-binding propensity

We assembled a dataset of experimentally validated mutations known to alter RNA-binding activity by querying UniProt entries using the search string ft_mutagen_exp:“RNA binding”. The retrieved records were pre-processed through a combination of automated cleaning and manual curation to standardize the *Mutagenesis* annotations. Because many UniProt entries report multiple mutations within a single annotation field, we parsed and split these entries such that each row in the final dataset corresponds to a single mutational event.

The curated dataset contains 393 mutations mapped to 136 proteins from 37 organisms. Of these, 295 mutations involve single–amino acid substitutions, whereas 98 correspond to multi–amino acid changes. The multi-residue mutations include 48 cases affecting adjacent residues and 50 involving non-adjacent positions. This curated set provides a reference collection for assessing how sequence variation modulates protein RNA-binding propensity.

#### Mutation score definition

To quantify mutation-associated changes in RNA-binding propensity, HERCULES computes residue-level RNA-binding propensity profiles for both the wild-type and mutant protein sequences. Let P^i^_WT_ and P^i^_MUT_ denote the predicted RNA-binding propensity at residue i in the wild-type and mutant sequence, respectively. The residue-wise profile difference is defined as:

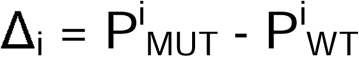

The primary output of HERCULES is the complete wild-type and mutant residue-level profiles, together with their residue-wise difference. For ranking and benchmarking mutations, we summarize the profile difference using a scalar mutation score defined as the difference between the mean mutant and wild-type profiles:

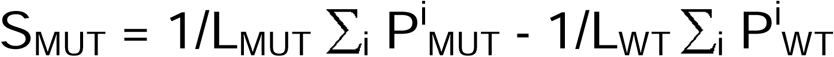

where L_WT_ and L_MUT_ are the lengths of the wild-type and mutant sequences. Negative values indicate a predicted decrease in RNA-binding propensity in the mutant sequence. This formulation is applicable to single amino-acid substitutions, multiple substitutions and length-altering variants such as insertions or deletions. The rationale for using a whole-profile summary, and its comparison with local mutation-centered indicators, is described in the **Supplementary Methods** and **Supplementary Fig. S16**. An auxiliary analysis testing whether HERCULES profiles encode quantitative information about experimental protein–RNA binding free-energy changes is described in the **Supplementary Methods**.

### Model architecture

#### Protein language model component

The protein language model component of HERCULES was based on ProteinBERT (23), a pretrained protein language model that combines local and global representations and can be fine-tuned for protein-level or residue-level prediction tasks. We fine-tuned ProteinBERT on a binary RNA-binding protein classification task using the implementation provided by the original ProteinBERT repository (https://github.com/nadavbra/protein_bert.git).

The benchmark dataset used for fine-tuning is described in the **Datasets** section. Protein sequences and binary labels were loaded from the corresponding training and test files. The original training file was then split into training and validation sets using a stratified random split, preserving the proportion of RNA-binding and non-RNA-binding proteins in both subsets. Specifically, 10% of the original training records were assigned to the validation set. This resulted in 3852 training records, 428 validation records, and 1070 independent test records. The independent test set was kept separate and was used only for final model evaluation, not for model selection or early stopping.

ProteinBERT was loaded using the “load_pretrained_model ()” function from the “proteinbert” Python package, which returns the pretrained model generator and the corresponding input encoder. The downstream prediction task was defined as a binary output with class labels 0 and 1 using “OutputType (False, “binary”)”. A fine-tuning model was then generated with “FinetuningModelGenerator”, using the binary output specification.

Fine-tuning was performed using the “finetune ()” function. Training was carried out with an initial sequence length of 512 amino acids, a batch size of 32, and a learning rate of 1 × 10−4. The pretrained layers were initially kept frozen, and the newly initialized task-specific layers were trained with a learning rate of 1 × 10−2. The maximum number of epochs per fine-tuning stage was set to 40. A final fine-tuning stage was then performed using a maximum sequence length of 1,024 amino acids, a learning rate of 1 × 10−5, and one final epoch.

Model optimization was monitored on the validation set. We used two Keras callbacks to control training. First, the learning rate was reduced when validation performance reached a plateau using “ReduceLROnPlateau”, with patience of 1 epoch, a reduction factor of 0.25, and a minimum learning rate of 1 × 10−5. Second, early stopping was applied using “EarlyStopping” with patience of 2 epochs and “restore_best_weights=True”, so that the final model corresponded to the epoch with the best validation performance before convergence. The independent test set was used only after training to assess final model performance.

After fine-tuning, residue-level profiles were derived from the attention values of the fine-tuned ProteinBERT model. To access the internal attention outputs, we used the ProteinBERT utility “get_model_with_hidden_layers_as_outputs”, which exposes hidden-layer outputs in addition to the final classification output. For each protein sequence, attention values were extracted across the model layers and heads and averaged to obtain a single per-residue ProteinBERT attention profile. This attention-derived profile was interpreted as the PLM-based contribution of each residue to the RNA-binding classification task. In the HERCULES framework, this profile is subsequently length-scaled and fused with the physicochemical profile to generate the final residue-level RNA-binding propensity profile.

#### Physicochemical descriptor-based model

The physicochemical component of the HERCULES framework is based on residue-level descriptors that capture local biochemical properties of protein sequences. For each wild-type (WT) and mutant protein in the curated datasets described above, we computed a total of 82 physicochemical features at the residue level, derived from established descriptor sets including the cleverSuite (40) and catGRANULE 2.0 ROBOT (25). These features encode diverse properties such as hydrophobicity, charge, flexibility, and structural propensity.

To identify the most informative physicochemical signals associated with loss of RNA-binding upon mutation, we performed feature selection using an elastic net regularized logistic regression model. The model was trained on paired wild-type (WT)–mutant differences, constructed such that positive examples corresponded to the correct direction of the physicochemical change (mutant minus WT), while negative examples were generated by reversing the sign of the difference vector. The AUROC of this model after feature selection is 0.93.

This formulation enforces sensitivity to the directionality of mutational effects and enables the model to distinguish meaningful physicochemical alterations from background variability. The elastic net regularization promotes sparsity while preserving correlated features, resulting in a robust and interpretable subset of physicochemical descriptors.

The feature selection procedure retained 30 physicochemical descriptors, effectively filtering out approximately 60% of the original feature pool. The selected features include properties such as hydrophobicity, charge, and intrinsic disorder, which are well established as key determinants of RNA-binding propensity in proteins. The enrichment of these biologically relevant descriptors among the selected features indicates that the model captures fundamental physicochemical principles underlying RNA–protein interactions. A detailed list of the retained features and their corresponding weights is provided in **Supplementary Fig. S1-S2**.

Using the selected descriptors, we derived an optimal linear combination that summarizes mutation-induced physicochemical changes into a single physicochemical profile. WT proteins and their corresponding mutants were encoded using the same physicochemical feature set, assuming a one-to-one pairing between sequences. For each pair, a difference vector was computed to isolate mutation-specific effects and remove background sequence variability. To derive the optimal weights we perform a classical procedure defined as Linear Discriminant Analysis defined by Fisher (41).

Let 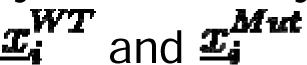 denote the physicochemical feature vectors of the i-th WT sequence and its mutant counterpart. For each pair, the difference vector is defined as

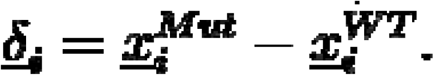

The mean difference vector ***<u>μ</u>*** and covariance matrix **Σ** are estimated across all n pairs. The optimal weight vector is ***<u>w</u>*** obtained by maximizing the signal-to-noise ratio

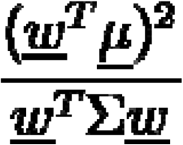

yielding the closed-form solution

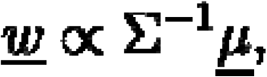

up to a scaling factor. Each mutation is assigned a scalar score

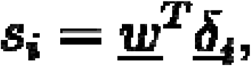

which quantifies the magnitude and direction of the physicochemical change induced by mutation. The method relies on the assumption that each feature follows a Gaussian distribution, which is satisfied in our data (**Supplementary Fig. S3**). The learned feature weights are reported as bar plots in **Supplementary Fig. S1-S2**, quantifying the relative contribution of each physicochemical descriptor to mutation sensitivity. Notably, individual physicochemical features already exhibit some discriminatory power in separating deleterious RNA-binding mutations within the corresponding WT sequence (**Supplementary Fig. S3**). However, the optimized linear combination of features substantially enhances performance, yielding an improvement of more than 20% compared to any single descriptor alone.

#### Profile fusion and weighting strategy

To integrate physicochemical information with contextual sequence representations, the physicochemical profile was combined with attention-based profiles derived from the fine-tuned ProteinBERT model. Because attention values inherently scale with protein length, the attention profiles were rescaled by the sequence length to ensure comparability across proteins of different sizes.

The final residue-level profile was obtained by combining the rescaled attention signal with a smoothed version of the physicochemical profile. Smoothing was performed using a sliding window of size *l*, allowing the physicochemical signal to capture broader sequence context. The two components were then integrated through a weighted combination, where the parameter α controls the relative contribution of the physicochemical signal.

The parameters governing smoothing and weighting were optimized to maximize the separation between high- and low-scoring regions within Pfam-annotated domains and to improve the identification of deleterious mutations (see “**Datasets**” for details). This procedure yielded an optimal window size equal to the protein length, corresponding to global averaging of the physicochemical signal, and a weighting factor α = 0.2. This strategy ensures that the final profile captures both contextual information from the language model and mutation-sensitive physicochemical effects. For visualization and web-server display, HERCULES residue-level profiles were placed on a common scale using fixed z-score calibration parameters estimated from the union of the training and test sets. This display normalization was not used for model training, parameter selection, threshold optimization, mutation-score computation or quantitative benchmarking. Specifically, the mean of the per-protein profile means and the mean of the per-protein profile standard deviations were estimated and used to perform z-score normalization of the fused profiles. The resulting output is a normalized residue-level RNA-binding propensity profile, hereafter referred to as the HERCULES profile.

#### Global score integration

In addition to residue-level predictions, HERCULES produces a global RNA-binding propensity score for each protein. Two complementary global scores were considered as inputs: (i) the global output score provided by the fine-tuned ProteinBERT model and (ii) the average value of the combined physicochemical profile across residues. These scores were integrated using a multilayer perceptron (MLP) trained on the same balanced dataset previously employed for binary classification of RNA-binding proteins.

The neural network learns a nonlinear combination of the sequence-based and physicochemical signals, yielding a fused global RNA-binding propensity score, referred to as the HERCULES score.

### Model validation and performance

Performance metrics, benchmarking strategies against existing methods and HERCULES model ablation analyses are reported in the **Supplementary Methods**.

### PDB protein–RNA complex analysis and exploratory AlphaFold3 augmentation

Experimentally resolved protein–RNA complexes were collected from the PDB as described in the Datasets section. Residue-level RNA-contact labels were assigned from experimental structures using a 7 Å distance cutoff between protein and RNA atoms. This benchmark was used as a complementary analysis because each PDB structure captures contacts with one RNA molecule in one structural context and therefore provides incomplete labels for general RNA-binding propensity.

For exploratory contact augmentation, a subset of proteins was modelled with AlphaFold3 in complex with a panel of structured G-quadruplex RNAs. AlphaFold3-derived contacts were not used as definitive ground truth, but only to expand the diversity of RNA-binding contexts. Models were retained based on predicted confidence scores, and RNA-contact masks were generated using the same 7 Å distance cutoff. Experimental and predicted masks were combined to form augmented annotations. Performance was evaluated using per-protein top-versus-bottom AUROC. Full details of the RNA panel, confidence filtering, contact-mask construction, dataset characterization and sequence-redundancy analyses are provided in **Supplementary Methods** and **Supplementary Figs. S8–S11**.

#### Protein array validation

RNA-binding was experimentally evaluated using the Cambridge Protein Arrays HuProt™ Human Protein Microarray v4.0, comprising over 20,000 individually printed recombinant full-length human proteins representing more than 16,000 human genes. Proteins were expressed as GST fusion proteins in *Saccharomyces cerevisiae*, purified, and immobilized in duplicate on glass slides, providing broad coverage of the human proteome for high-throughput protein–RNA interaction screening.

Three synthetic RNA probes corresponding to disease-associated repeat expansion sequences were generated by in vitro transcription from DNA templates containing 99 CGG repeats (CGG), 40 interrupted CAACAG repeats (CAACAG), or 23 CAG repeats (CAG). Following transcription, residual DNA templates were removed by DNase treatment, and RNAs were purified using magnetic bead purification. RNA integrity was verified by agarose gel electrophoresis and Agilent Bioanalyzer analysis. Purified RNAs were fluorescently labelled with Cy5, and labelling efficiency was quantified spectrophotometrically before hybridization.

For each RNA probe, 50 pmol of Cy5-labelled RNA were hybridized independently to HuProt arrays in three technical replicates. Following hybridization, arrays were washed, scanned at 635 nm using an Agilent G2505B microarray scanner, and fluorescence intensities were extracted with GenePix Pro 6.1 software. Foreground fluorescence signals were corrected for local background, duplicate protein spots were averaged, and only reproducible signals across technical replicates were retained for downstream analysis.

Replicate fluorescence intensities were averaged for each protein and mapped to gene symbols and UniProt identifiers. Array positivity was defined independently for each RNA probe using a robust z-score approach. For each array, the median and median absolute deviation (MAD) of fluorescence signals across all spotted proteins were calculated, and robust z-scores were computed using the standard 1.4826 × MAD scaling factor. Proteins with robust z-score ≥ 3 were classified as positive. For the combined analysis, a protein was considered array-positive if it exceeded this threshold for at least one of the three RNA probes.

Computational predictions were evaluated against the experimental measurements using fixed decision thresholds applied uniformly across the human proteome (HERCULES global score ≥ 0.565 and HydRA score ≥ 0.893). Agreement between predicted RNA-binding proteins and experimentally identified array-positive proteins was quantified using the Jaccard index.

Residue-level HERCULES predictions were subsequently examined for representative candidate proteins identified by the arrays. Candidates were required to (i) be absent from the human RBP2GO reference catalogue, (ii) exceed the HERCULES global RNA-binding threshold (score ≥ 0.565), (iii) be positive for at least two of the three RNA probes (robust z-score ≥ 3), and (iv) display extensive residue-level RNA-binding propensity, defined as at least 50% of residues with positive HERCULES scores. Application of these criteria yielded 18 candidate proteins for detailed residue-level analysis.

## Results

### A sequence based framework combining a protein language model and physicochemical descriptors

We designed HERCULES to predict RNA-binding propensity from protein sequence (**Fig. 1**; **Methods**). The model integrates two complementary components: a fine-tuned protein language model (ProteinBERT) (23) and a physicochemical descriptor-based model (25, 40), which are subsequently combined to generate a global RNA-binding score, residue-level profiles and a mutation score. We first fine-tuned ProteinBERT on a curated dataset of human RNA-binding proteins (RBPs) versus non-RBPs (25, 28, 29, 33, 34), treating the task as binary classification (see **Methods** and **Supplementary Table S1**). This procedure produces a ProteinBERT global score for each protein, reflecting the model-derived probability that the input sequence belongs to the RBP class. Importantly, we found that averaging the attention heads after fine-tuning provides a residue-level RNA-binding profile that enriches high-scoring residues within annotated RNA-binding regions (see **Methods**). This approach leverages the intrinsic representations learned by ProteinBERT during fine-tuning, without requiring additional post-hoc analysis steps, such as occlusion-based strategies previously employed, for instance, by HydRA (11). Protein language models are known to excel at capturing global, long-range sequence dependencies and evolutionary constraints (21, 23, 42, 43), while being less directly sensitive to fine-grained local physicochemical perturbations, such as those induced by single-residue mutations. To complement the protein language model with information on the local sequence context, we therefore constructed a residue-level physicochemical model. We collected a dataset of experimentally validated mutations that impair RNA binding from the UniProt database (44) (see **Methods**). We computed 82 residue-level descriptors capturing physicochemical properties (25, 40), performed feature selection to identify the most informative descriptors, and trained a linear model to predict the effect of mutations (see **Methods**). This component provides mutation-sensitive scores that reflect how local chemical environments influence RNA binding.

**Figure 1.**
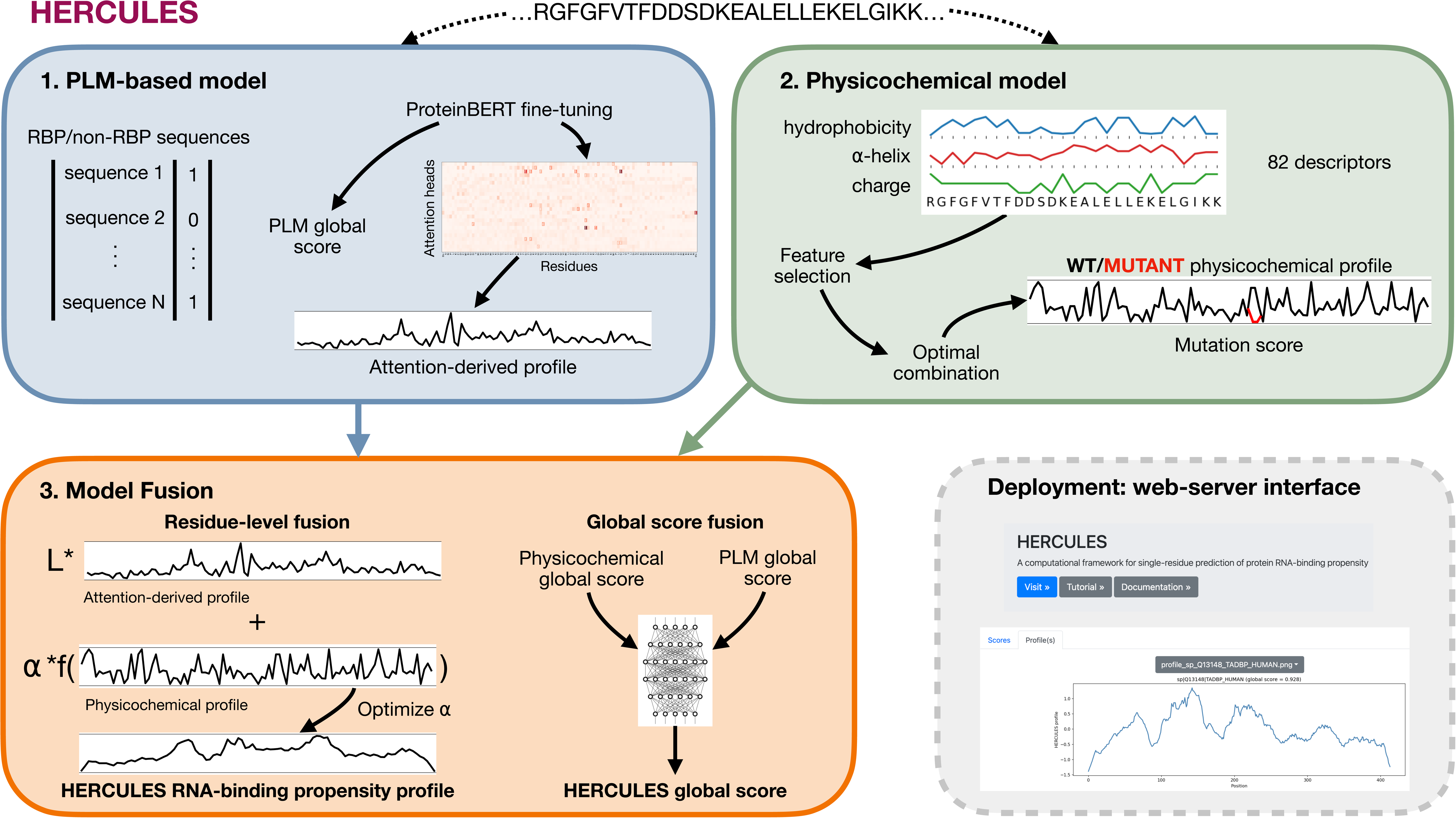
Overview of the HERCULES framework for sequence-based prediction of RNA-binding propensity. Schematic of the HERCULES architecture. A protein language model (PLM; ProteinBERT) is fine-tuned to discriminate RNA-binding proteins (RBPs) from non-RBPs, yielding a ProteinBERT global RNA-binding score and residue-level profiles obtained by averaging attention values across heads and layers. In parallel, a physicochemical model computes 82 residue-level descriptors, performs feature selection, and predicts mutation-sensitive effects based on experimentally validated RNA-binding–disrupting variants. The PLM-derived attention profile and the physicochemical profile are combined through optimized linear weighting to produce the final HERCULES residue-level RNA-binding propensity profile. Global scores from the PLM and the mean physicochemical profile are integrated by a multilayer perceptron to generate the final HERCULES global RNA-binding score. The HERCULES web server outputs include the residue-level profile, the RNA-binding global score and a mutagenesis heatmap summarizing mutation-associated changes in predicted RNA-binding propensity. Black arrows indicate within-module processing, colored arrows indicate module outputs entering the fusion step, and the dashed grey box denotes the deployment/web-server interface.

To combine the PLM-derived attention profile with the physicochemical profile, we performed a systematic search over a weighting parameter α using a dataset of human RBPs with annotated RBDs from the Pfam database (35), and a curated dataset of UniProt mutations experimentally shown to impair RNA binding, jointly optimizing for accurate identification of RBDs and for correct prediction of deleterious RNA-binding mutations (**Fig. 1** and **Supplementary Fig. S1-S3**; see **Methods**). This procedure identifies the optimal linear combination of the attention-derived and physicochemical profiles, resulting in a final residue-level RNA-binding propensity profile that is sensitive to both domain localization and mutation effects (11, 14–16). In addition to the residue-level profile, HERCULES reports a final global RNA-binding score, which is based on the ProteinBERT global score, defined as the output of the fine-tuned PLM classifier, and the physicochemical global score, defined as the mean of the physicochemical profile across residues. These two quantities are integrated using a multilayer perceptron trained on a balanced RBP versus non-RBP dataset, yielding the final HERCULES global score.

Evaluation of the HERCULES global score on an independent test set for discriminating RBPs yielded an area under the receiver operating characteristic curve (AUROC) of 0.86 (**Supplementary Fig. S4A**) and an area under the precision–recall curve (AUPRC) of 0.86 (**Supplementary Fig. S4B**).

Summing up, given an input protein sequence, HERCULES returns: (i) a global RNA-binding propensity score, (ii) a residue-level RNA-binding profile, and (iii) a mutation score for any given single- or multiple-residue variant relative to the wild-type sequence. This integrated framework enables prioritization of RNA-binding regions and mutation-associated propensity changes using only protein sequences.

### Residue-level localization of annotated RNA-binding regions

A central goal of HERCULES is the residue-level localization of RNA-binding regions along protein sequences, rather than the mere discrimination of RBPs from non-RBPs. To systematically evaluate residue-level performance, we benchmarked HERCULES against state-of-the-art residue-level predictors — HydRA (11), DRNApred (14), HybridRNAbind (16) and GPSite (17) — on a curated dataset of 604 human proteins with Pfam-annotated RBDs and sharing less than 30% sequence identity with the training set (**Fig. 2A–B**; see **Methods** and **Supplementary Table S2**). For each protein, we computed the AUPRC for discriminating residues within Pfam-annotated RBDs from residues outside annotated RBDs, and normalized it by the expected AUPRC of a random predictor, given by the fraction of annotated RBD residues in the sequence (see **Supplementary Methods**). As shown in **Fig. 2A**, HERCULES achieves a significantly higher AUPRC ratio to random than all competing methods, indicating a markedly improved ability to concentrate high scores on Pfam-annotated RNA-binding regions. We further assessed performance using the positive likelihood ratio (PLR) as a function of sensitivity, following the evaluation strategy originally introduced for HydRA (11). This metric quantifies how much more likely a residue predicted as RNA-binding is to be truly RNA-binding compared to a random residue. Across the full sensitivity range, HERCULES consistently exhibits higher PLR than all other methods (**Fig. 2B**), demonstrating superior enrichment of annotated RNA-binding residues among high-scoring positions. Together, these two complementary metrics indicate that HERCULES provides more informative domain-scale RNA-binding propensity profiles than existing approaches.

**Figure 2.**
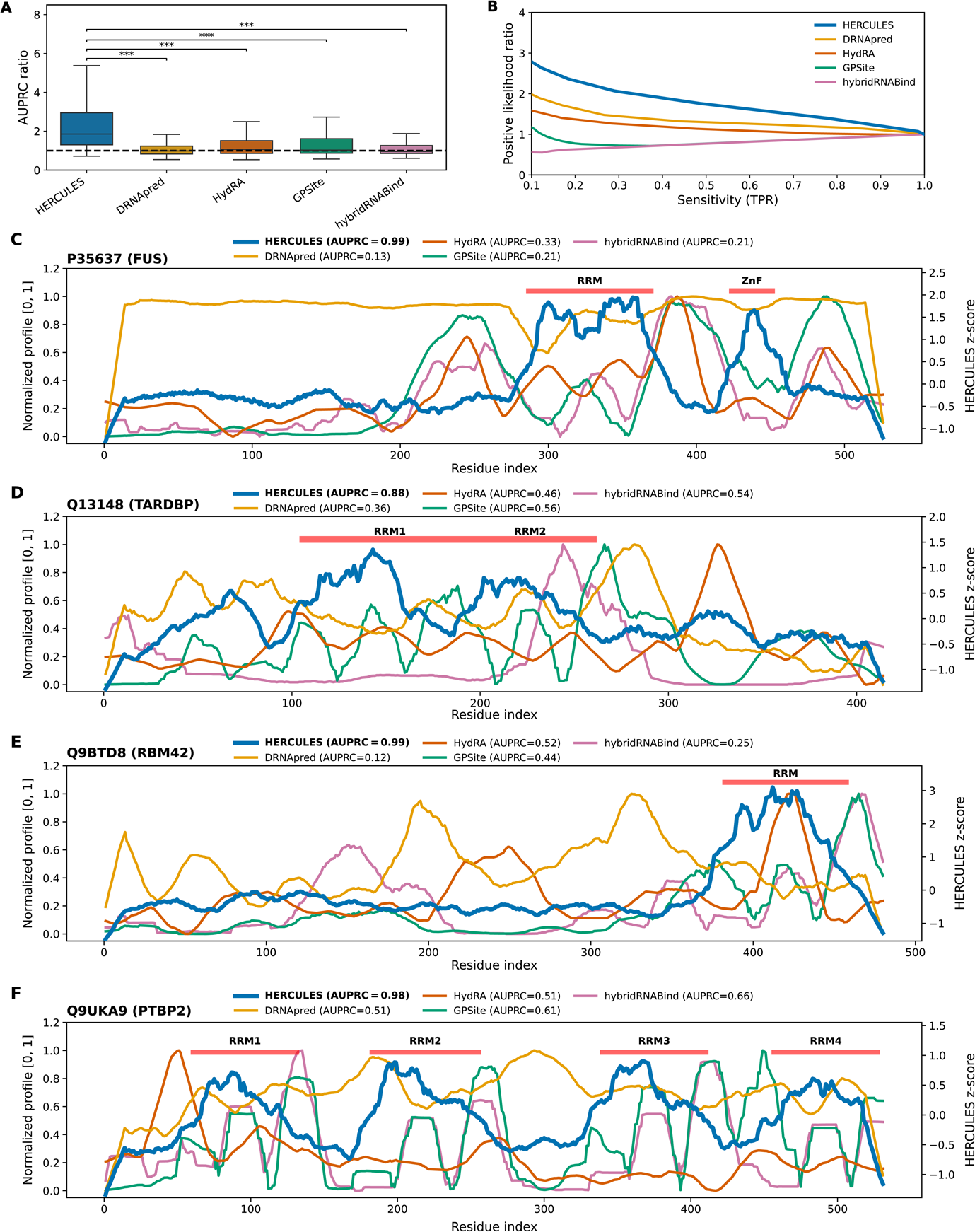
Validation of residue-level RNA-binding propensity profiles on Pfam-annotated RNA-binding regions. **(A)** Box plots showing the distribution of the Area Under the Precision–Recall Curve (AUPRC), normalized by the expected AUPRC of a random predictor (i.e., the fraction of RNA-binding residues in each protein), computed over 604 human proteins with Pfam-annotated RNA-binding domains and sharing less than 30% sequence identity with the training set. Colors indicate different prediction algorithms. Statistical significance versus HERCULES was assessed by paired Wilcoxon signed-rank tests on per-protein AUPRC ratios, with Bonferroni-adjusted p-values (***, padj < 0.001). **(B)** Positive likelihood ratio as a function of sensitivity for the same protein set shown in panel A. **(C–F)** RNA-binding propensity profiles for representative RNA-binding proteins containing distinct RNA-binding domains. Colored curves correspond to predictions from different algorithms, while annotated RNA-binding domains are indicated by red horizontal bars.

The practical impact of this improvement is illustrated by representative RNA-binding propensity profiles for well-characterized RBPs containing distinct RNA-binding domains (**Fig. 2C–F**). For FUS, which harbors an RRM and a zinc finger domain (26) (**Fig. 2C**), HERCULES highlights both annotated domains, whereas alternative methods produce weaker, broader, or more fragmented profiles. A similar pattern is observed for TDP-43 (**Fig. 2D**), RBM42 (**Fig. 2E**) and PTBP2 (**Fig. 2F**), where HERCULES more consistently resolves the annotated RRM regions across the full domain extent. These examples highlight that the gains observed at the population level translate into qualitatively improved domain-level annotations for individual proteins.

In **Supplementary Fig. S4A–B**, we show that, when evaluated strictly as a binary classifier of RBPs versus non-RBPs on the independent test set filtered at <30% sequence identity to the training set, HydRA slightly outperforms HERCULES in terms of AUROC and AUPRC. When extending the comparison across nine organisms using RBPs curated from the RBP2GO database (30), performed after the same <30% sequence-identity filtering, binarized HERCULES scores consistently identify a larger fraction of annotated RBPs than HydRA (**Supplementary Figure S4C; see Methods**). These analyses evaluate complementary properties. The independent human test set measures binary discrimination between RBPs and a defined negative set, whereas the RBP2GO analysis measures recovery of curated positive RBPs across heterogeneous organisms. Thus, HydRA performs better as a strict binary classifier, while HERCULES shows broader recovery of annotated RBP2GO positives after homology filtering, consistent with its intended use for detecting proteins that may contain localized RNA-binding regions. Notably, despite the fact that the residue-level profile is not constructed to encode global information, the HERCULES global score and the mean of the residue-level profile display a substantial correlation (Spearman ρ ≈ 0.6; **Supplementary Fig. S4D**). Consistently, the global score robustly separates RBPs from non-RBPs with a very large effect size (Cliff’s delta = 0.79; **Supplementary Fig. S4E**), while the mean of the residue-level profile also achieves significant discrimination, albeit with lower effect size (Cliff’s delta = 0.45; **Supplementary Fig. S4F**). Together, these observations show that, although HERCULES explicitly decouples global and local predictions by design, the residue-level profiles retain meaningful global information, reinforcing the biological consistency of the framework without compromising its primary objective of RNA-binding region localization. Ablation of the global score showed that the fused HERCULES global score and the ProteinBERT global score achieved essentially identical AUROC values on the independent test set (**Supplementary Fig. S5A–B** and **Supplementary Methods**), which indicates that the binary RBP classification is driven by the fine-tuned PLM component.

Because proteins differ substantially in length and in the fraction of residues annotated as RNA-binding, raw AUPRC values have different random baselines across proteins. We therefore used the ratio between the AUPRC and the protein-specific random baseline as the primary per-protein enrichment metric. However, additional analyses further support the robustness of HERCULES at the residue level (**Supplementary Fig. S6**). First, HERCULES also achieves the highest distribution of raw AUPRC values across the Pfam RBD benchmark, indicating that the improvement is not an artifact of normalization by the random baseline (**Supplementary Fig. S6A**). Second, when progressively comparing the highest- and lowest-scoring residues within each protein, HERCULES reaches the highest top-versus-bottom AUROC among the evaluated methods (**Supplementary Fig. S6B**). Third, HERCULES shows the strongest enrichment of annotated RNA-binding residues among the top 10% highest-scoring positions (**Supplementary Fig. S6D**), and achieves the best Matthews correlation coefficient distribution (**Supplementary Fig. S6E**; see **Supplementary Methods**). Together, these complementary metrics confirm that HERCULES concentrates high residue-level scores within annotated RNA-binding regions more effectively than the other methods. Notably, when stratifying proteins by their overall intrinsic disorder content, HERCULES performance in identifying RNA-binding domains improves with increasing disorder (**Supplementary Fig. S6C**). This behavior is consistent with the known enrichment of RNA-binding functions in intrinsically disordered regions (45) and suggests that HERCULES effectively exploits sequence features characteristic of disordered, interaction-prone segments. In contrast, the other evaluated methods remain substantially less enriched on Pfam-scale RBD annotations across disorder bins, indicating that contact-oriented or occlusion-based profiles are less aligned with domain-scale annotations in highly disordered proteins. These results indicate that HERCULES is particularly suited to capture RNA-binding determinants in disordered proteins, which represent a large and functionally important fraction of the RBP repertoire.

We further constructed an independent InterPro (46) member-database benchmark using RNA-binding region annotations supported by PROSITE (47), SMART (48) and not present in Pfam (**Supplementary Table S3**). After applying the same <30% sequence-identity filter against the HERCULES training set, the final benchmark contained 140 proteins. HERCULES retained the strongest enrichment of high-scoring residues within annotated RNA-binding regions across different performance metrics, supporting the robustness of the residue-level profiles beyond Pfam-derived annotations (**Supplementary Figure S7**).

As an additional exploratory analysis, we examined residue-level predictions on experimentally resolved protein–RNA complexes from the Protein Data Bank (PDB). This benchmark differs substantially from the Pfam and InterPro RBD benchmarks: PDB structures are typically shorter protein fragments or isolated domains, contain a smaller fraction of annotated RNA-contacting residues, and label only residues contacting the specific RNA molecule present in the solved complex (**Supplementary Fig. S8**). Thus, single-complex contact maps provide high-confidence but incomplete annotations for evaluating general RNA-binding propensity profiles. To assess how this limitation affects residue-level evaluation, we generated supplementary AlphaFold3 protein–RNA models for a subset of proteins using a panel of structured G-quadruplex RNAs (see **Methods, Supplementary Fig. S9** and **Supplementary Methods**). Indeed, G-quadruplexes are promiscuous RNA motifs known to engage protein regions beyond classically defined RNA-binding domains (27, 49, 50). These predictions were not treated as definitive ground truth, but only as exploratory structural augmentation to increase the diversity of RNA-binding contexts. In this analysis, HERCULES showed an increase in median top-versus-bottom AUROC, whereas DRNApred and HydRA decreased (**Supplementary Fig. S10**). This result suggests that some high-scoring HERCULES regions may correspond to RNA-interaction sites not observed in a single experimental complex. Because several PDB-trained predictors have substantial overlap with the proteins in this structural benchmark, we also quantified sequence similarity to available training sets. NCBRPred showed near-complete redundancy with the test proteins and was therefore excluded from the primary comparison, whereas DRNApred showed lower overlap (**Supplementary Fig. S11**).

To determine which model component drives residue-level localization, we performed a profile-level ablation on the Pfam benchmark dataset. The ProteinBERT attention-derived profile and the final HERCULES profile showed similar AUPRC/random distributions (**Supplementary Fig. S5C**), indicating that RBD-scale localization is primarily encoded by the PLM-derived attention profile, while the physicochemical component acts as a local modulation rather than as a standalone domain-localization predictor.

Overall, these results demonstrate that HERCULES substantially advances the state of the art in residue-level prediction of RNA-binding domains, providing accurate, well-localized profiles that outperform existing methods across diverse metrics and protein contexts.

### Identification of canonical, non-canonical and putative RNA-binding domains

To assess the ability of HERCULES to identify different classes of RBDs, we evaluated residue-level predictions on a comprehensive dataset integrating canonical, non-canonical, and putative RBD annotations derived from the Pfam database (9, 35) (see **Methods** and **Supplementary Table S2**). After filtering against the HERCULES training set, the final non-redundant benchmark comprised 604 human proteins, corresponding to 643 protein–RBD-class entries: 215 proteins with canonical RBDs, 248 with non-canonical RBDs and 180 with putative RBDs (9).

We first compared HERCULES with DRNApred, HydRA, HybridRNAbind and GPSite, using the AUPRC ratio to random, computed separately for each domain class (**Fig. 3A; Methods**). HERCULES achieved the highest AUPRC ratio across all three classes, with statistically significant improvements over all competing methods except GPSite in the canonical RNA-binding class, where the difference favored HERCULES but was not significant. As expected, performance was highest for canonical RNA-binding domains, which include well-characterized motifs such as RRMs, dsRBDs, and KH domains, reflecting their strong and conserved sequence signatures. Performance decreased for non-canonical domains and is lowest for putative domains; however, even in this most challenging category, HERCULES remains significantly better than a random predictor and outperforms other methods. This trend reflects the increasing uncertainty and heterogeneity of RNA-binding mechanisms moving from canonical to putative domains, rather than a limitation specific to HERCULES. We further assessed enrichment of annotated RBD residues using the positive likelihood ratio (PLR) as a function of sensitivity (**Fig. 3B–D**). Across all three RBD classes, HERCULES shows the highest PLR over the whole sensitivity range, indicating that high-scoring HERCULES residues are more strongly enriched within annotated RBDs than those predicted by the other methods. The advantage was particularly evident for non-classical and putative RBDs, where several competing predictors remain close to random expectation, whereas HERCULES retained robust enrichment.

**Figure 3.**
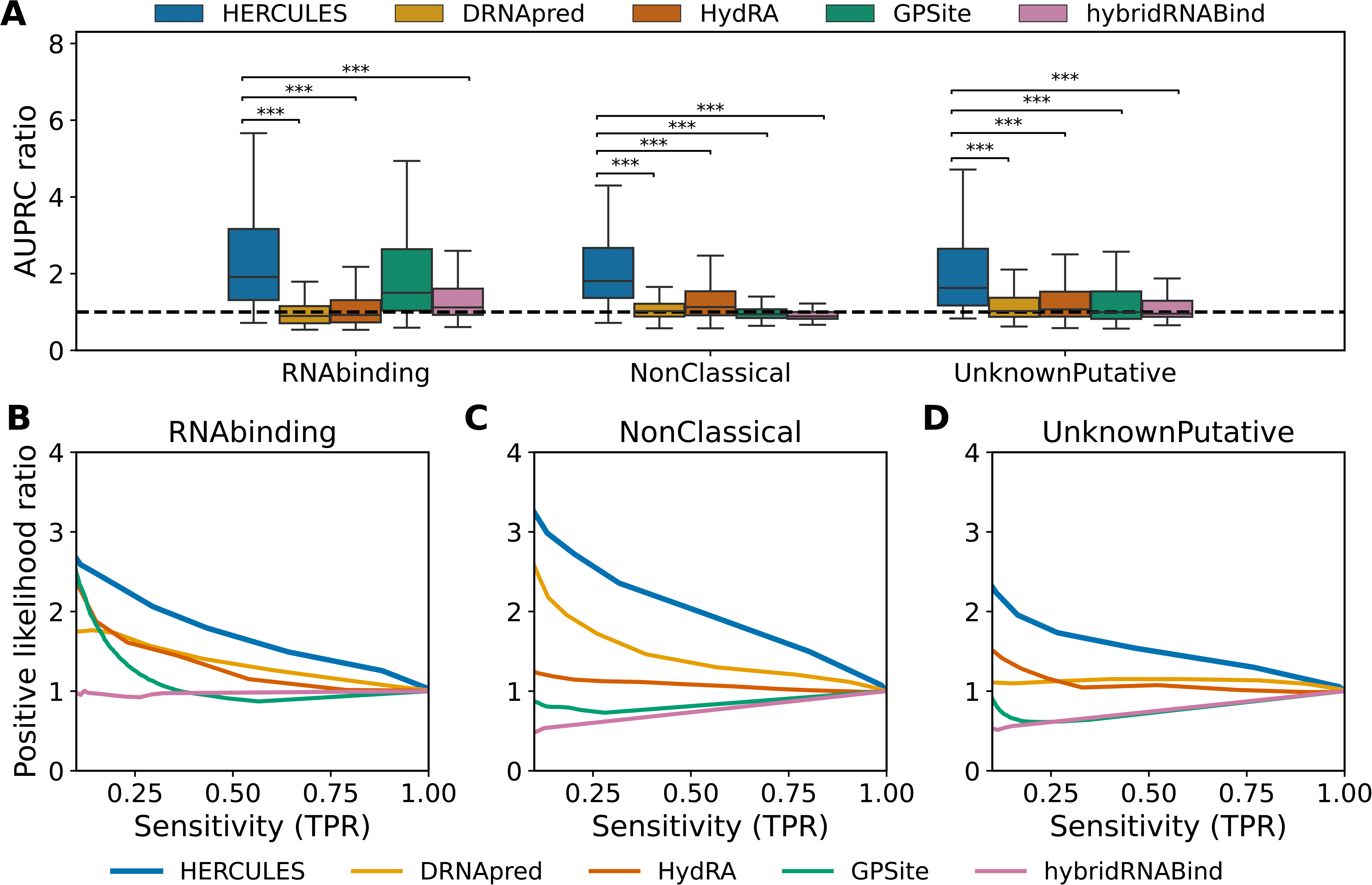
Recognition of canonical and non-canonical RNA-binding domains. **(A)** Box plots showing the distribution of the AUPRC ratio (AUPRC normalized by the expected value of a random predictor) for three classes of RNA-binding domains—RNABinding, NonClassical, and UnknownPutative—across five prediction algorithms (HERCULES, DRNApred, HydRA, GPSite and HybridRNAbind). Analyses were performed on Pfam-annotated human proteins filtered to <30% sequence identity with the HERCULES training set. Statistical significance versus HERCULES was assessed using paired Wilcoxon signed-rank tests on per-protein AUPRC ratios within each domain class, with Bonferroni correction for multiple pairwise comparisons. Asterisks denote Bonferroni-adjusted p-values: *** padj < 0.001. **(B–D)** Positive likelihood ratio as a function of sensitivity for each RNA-binding domain class, respectively, for the five algorithms shown in panel A.

Validation using complementary metrics confirmed these conclusions. Fold enrichment of annotated RNA-binding residues within the top 10% highest-scoring positions mirrored the results observed in **Fig. 3A**, with HERCULES showing the strongest enrichment across all domain classes (**Supplementary Fig. S12A**). Similarly, AUROC analysis based on progressively increasing subsets of top- and bottom-ranked residues showed that HERCULES achieved the best discrimination for canonical, non-classical and putative RBDs (**Supplementary Fig. S12B–D**). DRNApred was the closest competitor in several settings, particularly in the AUROC-based analysis, but HERCULES remained consistently higher across domain classes and ranking thresholds.

Overall, these results demonstrate that HERCULES robustly identifies RNA-binding domains across a continuum ranging from well-characterized canonical motifs to non-canonical or putative domains. The gradual decrease in performance from canonical to putative RBDs reflects the biological complexity and reduced annotation confidence of these classes. Importantly, under stringent homology filtering and across multiple complementary metrics, HERCULES remains above random expectation and outperforms sequence-based, occlusion-based, hybrid and structure-informed predictors, highlighting its ability to generate RBD-scale RNA-binding propensity profiles for both established and unconventional RNA-binding regions.

### Predicting the effect of mutations on RNA-binding propensity

We next assessed whether HERCULES can prioritize mutations that impair RNA-binding activity. We used a curated dataset of experimentally validated RNA-binding–disrupting variants derived from UniProt (44) , including both single and multiple amino-acid substitutions across multiple organisms and proteins (see **Methods** and **Supplementary Table S4**) . To avoid information leakage from multiple variants of the same protein, we split the dataset at the protein level while preserving organismal composition between training and test sets. The resulting mutation dataset and the distribution of variants across proteins, organisms and annotated binding regions are summarized in **Supplementary Fig. S13**.

For each wild-type (WT) and mutant protein pair, HERCULES computes residue-level RNA-binding propensity profiles. The default mutation score is defined as the difference between the mean mutant and WT profiles, so that negative values indicate an overall loss of predicted RNA-binding propensity in the mutant sequence (see **Methods**). Using this sequence-derived score, HERCULES correctly identified 85% of experimentally supported mutations in the held-out test set, outperforming DRNApred and HydRA using the same strategy for the computation of the mutation score (**Fig. 4A** and **Supplementary Fig. S14A**).

**Figure 4.**
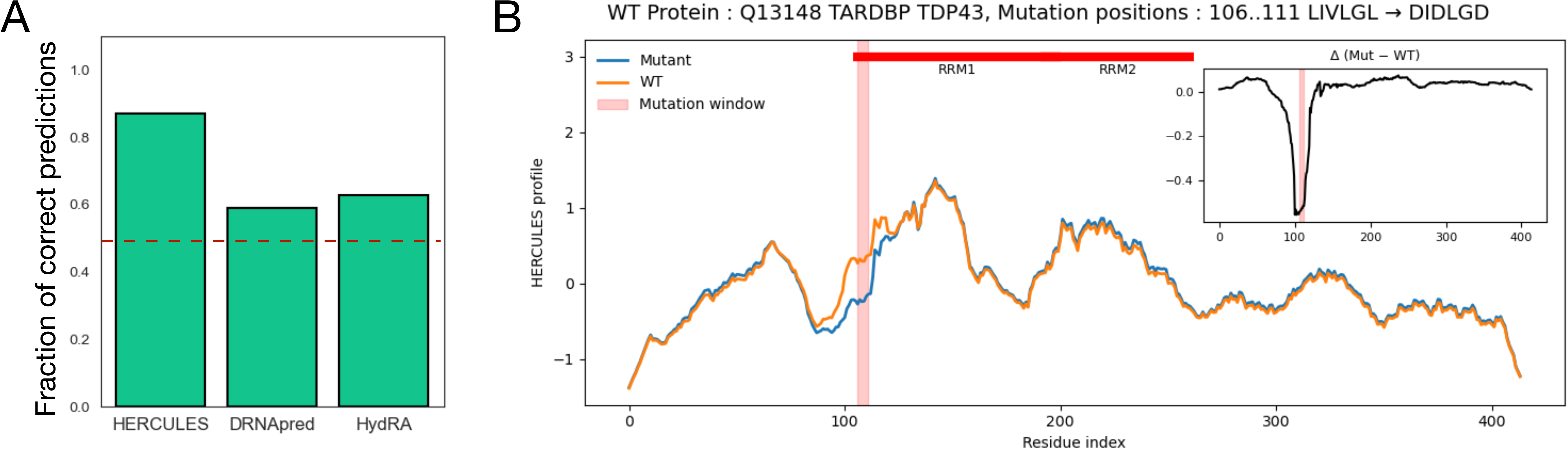
Prediction of the effects of mutations on RNA binding propensity. **(A)** Bar plot showing the fraction of mutations predicted as deleterious by HERCULES, DRNApred, and HydRA on the held-out test set. **(B)** HERCULES residue-level RNA-binding propensity profiles for the WT (orange) and mutant (blue) sequences around the mutation region. The example protein (TDP-43) carries a multiple amino-acid substitution (positions 106–111, LIVLGL → DIDLGD). The inset shows the profile difference (mutant − WT), highlighting a localized decrease in RNA-binding propensity at the mutation sites.

Although the mutation score is reported as a scalar for benchmarking and ranking, the primary output of HERCULES remains the complete WT and mutant residue-level profiles. These profiles allow the predicted effect to be localized along the sequence. For example, in TDP-43, a multiple amino-acid substitution produces a pronounced local decrease in predicted RNA-binding propensity near the mutated residues, while leaving much of the remaining profile largely unchanged (**Fig. 4B**). Additional representative variants show similar localized or regionally extended decreases in RNA-binding propensity around experimentally validated RNA-binding–abolishing mutations (**Supplementary Fig. S14C and Supplementary Fig. S15**). A mutation-centered heatmap of variants with the largest predicted effects further illustrates that the strongest WT–mutant differences often cluster around the mutated residues, while also revealing cases in which perturbations extend beyond the immediate local window (**Supplementary Fig. S15**).

We therefore compared local and global profile-difference indicators to determine how mutation effects are best summarized. Local windows around the mutated residues capture many short-range perturbations, but they do not fully account for variants that induce broader changes in the predicted RNA-binding profile. Consistently, the global mutant–WT mean difference showed stronger agreement with the experimental mutation labels than a strictly local score, with 88.2% of variants correctly assigned. Indeed, as shown in **Supplementary Fig. S16**, the fraction of correctly predicted mutations increases monotonically as the sequence window used to compute the mutation score is progressively enlarged around the mutation site. This supports the use of the global profile-difference score as a compact default summary, while retaining residue-level WT–mutant profiles for mechanistic interpretation. The same formulation also naturally extends to multi-residue substitutions and length-altering variants, for which a fixed local window may not fully capture the affected region.

We further investigated whether the HERCULES mutation score is associated with the structural environment of the mutated residue by analyzing its relationship with relative solvent accessibility (RSA) in the corresponding wild-type structure. RSA was computed as the solvent-accessible surface area normalized by the maximum accessible surface area expected for each amino-acid type. As shown in **Supplementary Fig. S17**, RSA is negatively correlated with the HERCULES mutation score (Spearman ρ = −0.403, p = 2.77 × 10−11), indicating that mutations at more solvent-exposed residues tend to produce larger predicted decreases in RNA-binding propensity. This trend is consistent with the predominantly surface-exposed nature of protein–RNA interaction interfaces and provides structural support for the interpretation that HERCULES prioritizes mutations affecting accessible residues likely to contribute to RNA binding.

We stress that the physicochemical component of HERCULES contributes most clearly in the mutation setting. A t-SNE projection of the selected physicochemical feature space separates wild-type and mutant sequences, indicating that the selected residue-level descriptors capture mutation-induced biochemical changes relevant to RNA binding (**Supplementary Fig. S14B**). Feature-family analysis further showed contributions from charge, disorder-related, hydrophobicity and secondary-structure-associated descriptors, consistent with known physicochemical determinants of protein–RNA interactions (**Supplementary Fig. S1** and **Supplementary Table S5**). In the ablation analysis, the physicochemical and fused HERCULES mutation scores identified a larger fraction of experimentally supported deleterious mutations than the ProteinBERT-only score, confirming that explicit physicochemical descriptors increase sensitivity to local amino-acid substitutions (**Supplementary Fig. S5D**).

Finally, we asked whether the full HERCULES residue-level profiles contain quantitative information about protein–RNA binding free-energy changes. Using the PRA-MutPred benchmark (51), we first compared the default HERCULES profile-difference score with experimental ΔΔG values. The score showed the strongest discrimination for mutations with the largest positive or negative ΔΔG values, whereas performance decreased when mutations with smaller energetic effects were included (**Supplementary Fig. S18A**). This indicates that the default mutation score captures large perturbations in RNA-binding energetics, but is not designed to resolve subtle quantitative affinity changes. We therefore trained a Gradient Boosting regression model to test whether additional ΔΔG information is encoded in the complete wild-type and mutant HERCULES profiles. The model was trained on the training split only, using wild-type and mutant HERCULES profiles as input features. The resulting predictions correlated strongly with experimental ΔΔG values across the benchmark (Pearson r = 0.861, Spearman ρ = 0.765; **Supplementary Fig. S18B**). This analysis indicates that quantitative energetic information is present in the HERCULES profiles, even though the default HERCULES output is intended for mutation-effect prioritization rather than direct ΔΔG prediction.

### Protein-array validation supports HERCULES-prioritized repeat-RNA-binding candidates

To provide orthogonal experimental support for HERCULES-prioritized RNA-binding candidates, we analyzed human protein arrays probed with three repeat-containing RNAs: CGG, CAG and CAACAG (see **Methods** and **Supplementary Table S7**). Among 14,064 proteins present on the arrays, 3,065 were array-positive using a robust threshold of z ≥ 3 in at least one probe, and 5,224 were classified as HERCULES-positive using the global-score threshold of 0.565. HERCULES-positive proteins showed consistent overlap with array-positive proteins across increasingly stringent signal thresholds, with Jaccard indices of 0.23, 0.18 and 0.16 for z ≥ 2, z ≥ 3 and z ≥ 4, respectively (**Supplementary Fig. S20A**). HydRA-positive proteins followed the same trend with somewhat lower Jaccard indices (0.17, 0.15 and 0.14), and both predictors showed overlaps of comparable statistical significance (hypergeometric p ≪ 0.001; **Supplementary Fig. S20A**).

We next focused on non-catalogued candidates by selecting proteins absent from RBP2GO that were HERCULES-positive, array-positive in at least two of the three probes, and displayed broad residue-level HERCULES signal (see **Methods**). This last criterion was included to prioritize proteins for which HERCULES did not merely assign a positive global score, but predicted extended RNA-binding propensity across a substantial fraction of the protein sequence. Thus, the analysis tests not only protein-level candidate prioritization, but also whether broad HERCULES residue-level profiles are supported by an independent RNA-binding readout. This filtering yielded 18 candidates (**Supplementary Fig. S20** and **Supplementary Table S8**).

Several candidates are zinc-finger or chromatin-associated proteins, including ZNF766, ZNF34, ZNF383, ZNF3, ZNF789, ZNF625, ZNF230, SPIN2B and PRM1. This composition is biologically plausible given recent systematic evidence that zinc-finger proteins, historically annotated mainly as DNA-binding transcriptional regulators, can also bind RNA and regulate post-transcriptional processes, including splicing, alternative polyadenylation, stability and translation (52). A complementary study using CLIP further reported widespread RNA binding among C2H2 zinc-finger transcription factors (53). The repeat-RNA probes used here are also expected to form structured RNA elements: CGG repeat RNA can adopt G-quadruplex conformations, and CGG-derived RNA G-quadruplexes have been shown to recruit protein interactors in fragile-X-related repeat-expansion contexts (54, 55). More generally, recent work identified nuclear RNA G-quadruplex-binding proteins, including many chromatin-associated proteins, supporting the idea that structured repeat RNAs can engage non-canonical RNA-binding proteins (50).

Representative examples further illustrate how array binding relates to the domain-scale HERCULES residue-level signal. SPIN2B was positive for all three repeat-RNA probes and displayed extended positive HERCULES regions that overlap its three Tudor domains (**Fig. 5A**). This correspondence is notable because, although Tudor domains are classically associated with recognition of methylated protein residues, direct RNA-binding activity has been demonstrated for members of the Tudor family. The Tudor domain of bacterial ProQ interacts with small non-coding RNAs, and the picocyanobacterial Tudor-domain protein PSHCP binds single-stranded, double-stranded and hairpin RNAs directly (56). Tudor-containing regions have also been implicated in RNA recognition by FMRP (9). Thus, despite SPIN2B not being annotated as an RBP in RBP2GO, the coincidence between array positivity, HERCULES-positive regions and Tudor domains suggests that these domains are plausible candidate RNA-interaction regions. Moreover, its homology to SPIN-family chromatin readers is notable because SPIN1 is a multivalent histone-mark reader implicated in chromatin regulation, rRNA transcription and piRNA-related biology (57, 58).A similarly striking correspondence is observed for ZNF625, where the oscillating HERCULES profile produces positive peaks across much of the tandem zinc-finger array (**Fig. 5B**). Classical C2H2 zinc fingers, traditionally regarded primarily as DNA-binding modules, are increasingly recognized as direct RNA-binding domains. For example, the C2H2 zinc fingers of YY1 bind single-stranded RNA, providing experimental evidence that this widespread domain architecture can mediate RNA recognition (59). The array positivity of ZNF625 for CAG and CAACAG RNAs, together with the correspondence between its zinc-finger architecture and HERCULES profile, is therefore consistent with an RNA-binding function distributed across its tandem zinc-finger region.

**Figure 5.**
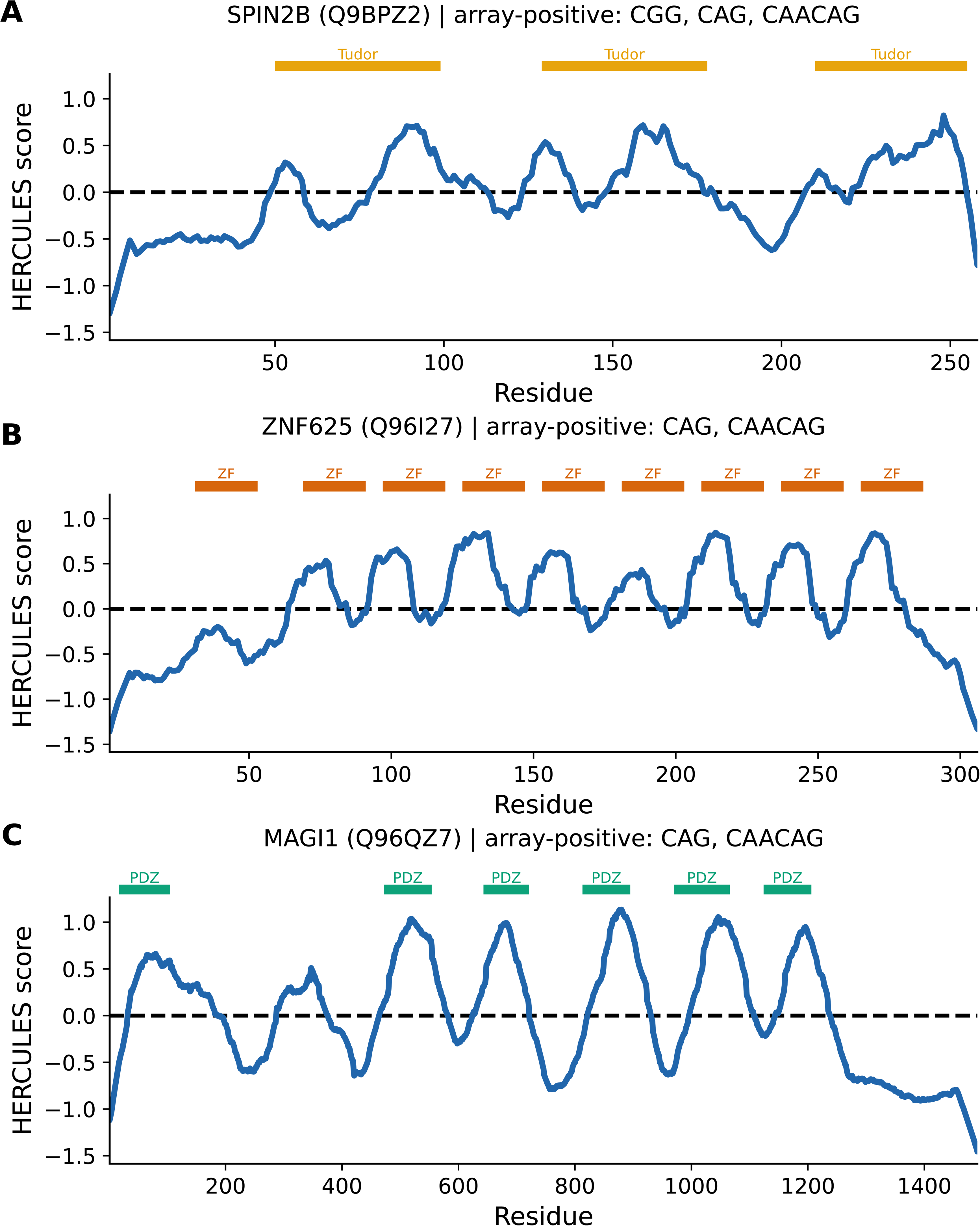
Protein-array validation of HERCULES predictions. Representative HERCULES residue-level profiles for three non-RBP2GO candidates that satisfy all selection criteria: SPIN2B (Q9BPZ2) **(A)**, ZNF625 (Q96I27) **(B)** and MAGI1 (Q96QZ7) **(C)**. Per-residue HERCULES scores are plotted along the sequence; the dashed line marks score = 0. Colored bars above each profile indicate UniProt-annotated domains (Tudor, orange; zinc finger, vermillion; PDZ, green). Panel titles report UniProt accessions and the subset probes on which each protein was array-positive (robust z ≥ 3). Full candidate set and selection criteria are provided in **Supplementary Figure S20** and **Supplementary Table S8**.

MAGI1 provides an especially interesting example of a less canonical RNA-binding architecture. Its strongest HERCULES peaks repeatedly coincide with its PDZ domains (**Fig. 5C**). Although PDZ domains are best known as protein–protein interaction modules, proteome-wide mapping of RNA-binding regions identified PDZ domains as direct RNA-binding modules and provided experimental validation of their RNA-binding activity (60). Thus, the recurrent array positivity of MAGI1 and the alignment of HERCULES maxima with multiple PDZ domains are consistent with an emerging non-canonical RNA-binding function for this domain family.

Collectively, these examples suggest that HERCULES does more than identify broadly RNA-binding proteins: in array-supported candidates, elevated residue-level propensity frequently maps onto recognizable domain architectures with independent experimental precedent for RNA binding. This agreement is particularly informative for non-canonical RNA-binding modules such as Tudor and PDZ domains and supports the use of HERCULES to prioritize specific domains for subsequent biochemical validation.

## Discussion

In this work, we introduced *cat*RAPID *signature* 2.0 HERCULES, a sequence-based framework for predicting RNA-binding propensity at both protein and residue resolution and for prioritizing sequence variants that may alter RNA-binding activity. HERCULES was designed to connect three tasks that are biologically related but often treated separately: global RBP classification, localization of RNA-binding regions and mutation-effect analysis. In this framework, a protein-level score summarizes global RNA-binding tendency, residue-level profiles identify regions contributing to this propensity, and wild-type versus mutant profile comparisons highlight sequence changes that may disrupt RNA binding. A central strength of HERCULES lies in its ability to generate interpretable residue-level profiles that reliably identify RNA-binding regions across diverse protein families. By leveraging attention patterns from a fine-tuned protein language model (23), HERCULES captures global sequence context, including domain architecture and long-range dependencies that are difficult to represent using local descriptors alone. These attention-derived profiles are particularly suited to RBD-scale prediction: they do not aim to reproduce only the sparse set of atomic contacts observed in one protein–RNA structure, but rather to highlight broader regions of the protein sequence that contribute to RNA-binding propensity. This distinction is important because functional RNA-binding regions can include canonical domains, disordered segments, non-canonical motifs or alternative interaction surfaces that may not be fully represented by a single structural complex.

Consistent with this interpretation, HERCULES enriches high-scoring residues within annotated RNA-binding regions across multiple benchmarks. On Pfam-annotated proteins (35), HERCULES outperforms sequence-based, hybrid and structure-informed algorithms, including DRNApred (14), HydRA (11), HybridRNAbind (16) and GPSite (17). The improvement is observed across complementary metrics and extends across canonical, non-classical and putative RNA-binding domain classes. The strongest performance is observed for canonical domains such as RRMs, KH domains and dsRBDs (35), where sequence signatures are better defined, but HERCULES also retains enrichment in less conventional RNA-binding regions (9, 36, 37). The independent InterPro benchmark (46) further indicates that the signal captured by HERCULES is not restricted to Pfam-derived annotations, supporting its use as a general RBD-scale RNA-binding propensity profiler.

A key contribution of HERCULES is the integration of residue-level RNA-binding propensity profiling with mutation-sensitive physicochemical modeling in a shared sequence-based framework. Deep neural network–based approaches, including AlphaFold (61) and AlphaFold3 (62), as well as large protein language models (21, 23, 42), have shown remarkable success in structure prediction and global functional inference. Nevertheless, the effect of individual amino-acid substitutions often depends on local biochemical changes, including charge, hydrophobicity, disorder propensity and secondary-structure preference. HERCULES addresses this point by incorporating an explicit physicochemical branch built from established residue-level descriptors (25, 40) and curated RNA-binding-disrupting mutations from UniProt (44). The ablation analyses support this division of labor: the ProteinBERT-derived branch provides the dominant signal for global RBP classification and RBD-scale localization, whereas the physicochemical component contributes most clearly to mutation sensitivity by locally modulating the PLM-derived profile.Although HERCULES is not designed as a dedicated binding-affinity predictor, the mutation profiles also contain quantitative information related to RNA-binding energetics. A supervised regression model trained on wild-type and mutant HERCULES profiles correlates with experimental ΔΔG values from the PRA-MutPred benchmark (51), indicating that the profiles encode biophysically meaningful information about mutation-induced affinity changes. This does not replace specialized ΔΔG predictors, which are trained specifically for affinity estimation and may incorporate structural information. Instead, it supports the interpretation that HERCULES profiles capture sequence-derived changes in RNA-binding propensity that are relevant for variant prioritization.

The biochemical experiments using protein arrays provide an independent experimental layer to this prioritization framework. HERCULES-positive proteins show reproducible overlap with proteins binding repeat-containing RNAs on human protein microarrays. We then asked whether HERCULES could prioritize proteins not currently annotated as RBPs. A small group of candidates was supported by a high HERCULES global score, binding to multiple repeat-RNA probes on the arrays, and extended positive HERCULES signal along the protein sequence. This is important because it connects protein-level prediction, residue-level profiling and experimental RNA-binding evidence in the same candidates. Several of these proteins belong to zinc-finger or chromatin-associated families, consistent with recent evidence that proteins traditionally annotated as DNA- or chromatin-associated can also interact with RNA and participate in post-transcriptional regulation (50). The repeat RNAs used in the arrays are also expected to form structured elements, which actively recruit protein interactors (4, 63). More generally, RNA G-quadruplexes engage diverse protein families (49, 50), supporting the plausibility that structured repeat RNAs can reveal non-canonical RNA-binding candidates (64). These array data therefore support HERCULES as a tool for candidate prioritization, while leaving the specificity and affinity of RNA targets to be further defined experimentally. Moreover, the AlphaFold3-augmented analysis (62) with structured G-quadruplex RNAs was used in this context to explore how additional RNA-binding contexts can alter contact annotations. The resulting observations are consistent with the idea that some HERCULES high-scoring regions may correspond to RNA-interaction surfaces not engaged in a single experimental complex. Thus, structural-complex analyses are best viewed as complementary to the Pfam, InterPro, mutation and array results, illustrating the relationship between context-specific contacts and domain-scale RNA-binding propensity.

Despite these advances, HERCULES also highlights current limitations of sequence-based RNA-binding prediction. The available experimental annotations are sufficient to robustly identify RNA-binding domains and mutation-sensitive regions, but remain too sparse to learn residue-level affinity and RNA-binding specificity across diverse RNA targets. Consequently, the mutation module relies on integrating protein language model representations with physicochemical descriptors to infer the local impact of amino acid substitutions, rather than directly learning binding energetics from experimental measurements. Future generations of HERCULES will exploit the rapidly expanding body of quantitative protein–RNA interaction data to move beyond domain localization toward prediction of RNA-binding affinity and sequence specificity. In this direction, HERCULES provides a key building block for the next generation of catRAPID (31, 38, 65), where robust identification of RNA-binding domains will be integrated with quantitative models of protein–RNA recognition. Together, these developments will enable increasingly accurate sequence-based prediction of protein–RNA interactions while preserving the interpretability and mutation sensitivity that characterize the present framework.

## Data availability

All datasets generated and analyzed in this study are provided as Supplementary Tables. HERCULES is freely available as an open-source Python package at https://github.com/tartaglialabIIT/hercules and through a public web server at https://tools.tartaglialab.com/hercules.

## Acknowledgments

The authors would like to thank the ‘RNA Flagship’ at IIT, Gabriele Proietti, and all the members of Tartaglia’s lab at IIT.

## Contributions

J.F. and M.M. developed and implemented the HERCULES methodology. G.G.T. conceived the model and supervised the study. D.M.V. evaluated the performance of the method across multiple datasets, including the AlphaFold3-based analyses. L.D.R. performed the analyses of experimentally resolved protein–RNA complexes from the Protein Data Bank. A.A. contributed to data analysis and developed the HERCULES web server. R.D.P. analyzed the protein-microarray data. V.V., T.B.O. and E.M. performed the protein-microarray experiments. J.F., M.M. and G.G.T. wrote the manuscript with contributions from all authors. All authors interpreted the results, reviewed and revised the manuscript, and approved the final version.

