## Supplementary Materials for "HERCULES: a deep-learning framework for RNA-binding region localization and mutation-effect prediction"

### Supplementary Methods

#### Global score

To assess the predictive performance of the HERCULES global score, we first evaluated it on an independent test set (see **Datasets** section in the **Methods**). To minimize overlap with model training, test proteins were screened against the HERCULES training set using MMseqs2 easy-search. Proteins were excluded if their best hit to a training sequence had sequence identity ≥30% and bidirectional alignment coverage ≥80%, where coverage was defined as min(query coverage, target coverage). This yielded a filtered independent test set of 979 proteins, comprising 481 RNA-binding proteins and 498 non-RNA-binding proteins; 91 proteins were removed from the original set of 1,070. Receiver operating characteristic (ROC) and precision–recall (PR) curves were computed from continuous global scores, and AUROC and AUPRC values were calculated using scikit-learn v1.3.2.

We further evaluated HERCULES on experimentally annotated RNA-binding proteins from RBP2GO across multiple organisms. The same MMseqs2 filtering criterion was applied to the RBP2GO reference set, retaining 108,248 of 132,205 screened proteins. Proteins whose UniProt identifiers were present in the HERCULES training set were also excluded. For organism-level benchmarking, we analyzed 12,472 RBP2GO-positive proteins from nine organisms: *Arabidopsis thaliana*, *Caenorhabditis elegans*, *Danio rerio*, *Drosophila melanogaster*, *Escherichia coli*, *Homo sapiens*, *Mus musculus*, *Plasmodium falciparum* and *Saccharomyces cerevisiae*. Continuous HERCULES scores were binarized using a threshold of 0.565, determined by maximizing Youden's J statistic on the validation set. The fraction of proteins predicted as RNA-binding was then computed for each organism.

For benchmarking, HERCULES was compared with HydRA. HydRA scores were generated using the HydRA command-line interface (HydRa2_predict; https://github.com/YeoLab/HydRA) and binarized using the threshold reported in the original HydRA manuscript (0.827)11. ROC and PR curves were computed for both methods on the independent binary-classification test set, and positive-prediction fractions were computed on the RBP2GO organism-level sets using the corresponding method-specific thresholds.

#### RNA-binding propensity profiles

We evaluated the ability of HERCULES RNA-binding propensity profiles to identify RNA-binding residues using a curated set of 604 human proteins with annotated RNA-binding domains (RBDs) from the Pfam database, with <30% sequence identity with proteins belonging to the training set (see the **Datasets** section in the **Methods**). Performance was assessed at residue resolution using multiple complementary metrics designed to capture different aspects of predictive quality.

#### Protein-Specific Enrichment of Precision–Recall Signal

For each protein and each algorithm, we computed the area under the precision–recall curve (AUPRC) using residue-level predictions and binary RBD annotations. To account for class imbalance and protein-specific RBD coverage, the AUPRC was normalized by the expected AUPRC of a random classifier, defined as the fraction of RNA-binding residues in the protein ($n_{pos}/L$, where $n_{pos}$ is the number of annotated RNA-binding residues and L is the protein length). This yielded an AUPRC ratio to random, with values >1 indicating enrichment over random expectation.

Pairwise comparisons between algorithms were performed using the Wilcoxon signed-rank test on per-protein AUPRC ratios, and effect sizes were quantified using rank-biserial correlation. Multiple testing correction was applied using the Bonferroni method.

#### Enrichment of top-ranked residues

To assess the ability of each method to prioritize RNA-binding residues, we computed the fold enrichment of annotated RNA-binding residues among the top 10% of residues ranked by predicted score. For each protein, residues were sorted by decreasing predicted propensity, and the fraction of true RNA-binding residues within the top-ranked subset was compared to the expected fraction under random selection ($n_{pos}/L$). Fold enrichment values >1 indicate preferential concentration of RNA-binding residues among top predictions.

#### Positive likelihood ratio

To evaluate the trade-off between sensitivity and specificity across score thresholds, we computed the positive likelihood ratio ($LR^+$) at residue level, previously employed by HydRA[1]. For a given algorithm and threshold t, residues with scores $\geq t$ were classified as RNA-binding. $LR^+$ was defined as:

$$\mathrm{LR}^+(t) = \frac{\mathrm{Sensitivity}(t)}{\mathrm{False\ Positive\ Rate}(t)} = \frac{\mathrm{TP/P}}{\mathrm{FP/N}}$$

where TP and FP denote the number of true and false positive residues, respectively, and P and N denote the total number of RNA-binding and non-RNA-binding residues. $LR^+$ was computed over 100 evenly spaced thresholds between the minimum and maximum predicted scores for each algorithm. When the false positive rate was zero, $LR^+$ was set to infinity if sensitivity was non-zero, and to zero otherwise. This metric quantifies the increase in odds of correctly identifying RNA-binding residues relative to misclassifying non-binding residues.

#### AUROC of top- and bottom-ranked residues

For increasing values of k, we selected the top k and bottom k residues ranked by predicted score and computed the AUROC distinguishing annotated RNA-binding residues from non-binding residues within this restricted set, as in a previous work[2]. This analysis evaluates how rapidly predictive signal concentrates at the extremes of the score distribution.

#### Benchmarking against existing methods

HERCULES was compared with several state-of-the-art predictors in residue-level and mutation-effect analyses. DRNApred and HydRA were included in both RBD localization and mutation-effect benchmarks. HybridRNAbind and GPSite were included in the RBD localization benchmark using their RNA-binding residue propensity outputs. In principle, HybridRNAbind and GPSite can also be evaluated for mutation effects by applying the same wild-type versus mutant profile-difference strategy; however, this analysis was not included here because mutation-effect scoring is not part of their original formulations and would require an additional post hoc benchmarking setup. HydRA residue-level profiles were derived using the occlusion_map3 command-line tool, inverted to match score directionality, padded to account for boundary effects introduced by occlusion, and normalized to the [0,1] range. HybridRNAbind was included as a sequence-based predictor of RNA-interacting residues across structure-annotated and disorder-annotated proteins. GPSite was included as a sequence-derived, structure-informed predictor of RNA-binding residues, using its RNA-binding output. For all methods, residue-level scores were normalized to the [0,1] range before comparison. NCBRPred was evaluated where computational constraints allowed, specifically on the protein–RNA complex dataset, given the web server submission limit of 10 sequences per job and intermittent server availability. NCBRPred is an extension of DRNApred.

#### Model ablation analysis

To quantify the contribution of each HERCULES component, we evaluated three model variants at global, residue and mutation level:

- **ProteinBERT**: fine-tuned ProteinBERT output alone (global score or L-scaled mean attention profile, where L is sequence length).
- **Physicochemical**: elastic-net weighted combination of amino-acid property scales (global mean or per-residue profile; weight $\alpha = 0.2$).
- **Fused (HERCULES)**: full model combining both branches (MLP fusion for global scores; $L \times$ attention + $\alpha \times$ mean(physchem) for profiles).

Physicochemical features were selected and weighted by elastic-net regression trained on the HERCULES training set. Residue-level profiles were smoothed (5% sequence-length moving average) and z-normalized using fixed calibration parameters ($\mu = 1.048$, $\sigma = 0.474$).

Global ablation was performed on an independent test set of 979 proteins (<30% sequence identity to the HERCULES training set by MMseqs2 easy-search; minimum alignment coverage 80%), comprising 481 RBP and 498 non-RBP proteins. Performance was assessed by area under the ROC curve (AUROC). Score distributions for RBP and non-RBP proteins were compared with two-sided Mann–Whitney U tests.

Residue-level ablation used the retained Pfam validation set (604 proteins, same <30% identity criterion). For each protein, three profile variants were computed and evaluated for RNA-binding domain (RBD) localization using the per-protein AUPRC ratio (model AUPRC divided by random expectation), as in the main Pfam benchmark.

Mutation ablation used experimentally annotated mutations from UniProt, split by protein into training (108 proteins) and test (28 proteins; 92 mutations) sets so that all mutations from a given protein belonged to the same partition. For each variant, wild-type and mutant sequences were scored and the mutation effect was defined as:

mutation score = mean(mutant profile) − mean(wild-type profile)

A negative score indicates predicted loss of RNA-binding propensity. Performance was summarized as the fraction of mutations with negative scores (“fraction of correct predictions”), under the expectation that experimentally disruptive mutations reduce global RNA-binding propensity.

**Mutation score evaluation and local-versus-global profile summaries**

The HERCULES mutation score is intended to summarize mutation-associated changes in RNA-binding propensity while preserving the complete wild-type and mutant residue-level profiles as the primary mechanistic output. Because the effect of a mutation may be local, regionally extended or distributed across multiple parts of a protein, we evaluated several profile-difference summaries before selecting the default mutation score.

For each wild-type and mutant sequence pair, HERCULES computes residue-level profiles $P^{WT}$ and $P^{MUT}$, and the residue-wise difference $\Delta_i = P^i_{MUT} - P^i_{WT}$. Negative values indicate lower predicted RNA-binding propensity in the mutant sequence. We then compared mutation scores computed over sequence windows centered on the mutated residue or residues. Window sizes ranged from local mutation-centered regions to the complete protein sequence. For each window, the mutation score was defined as the average mutant–wild-type profile difference within the selected region.

This analysis showed that the fraction of experimentally validated RNA-binding-disrupting mutations assigned a negative score increased as larger regions of the profile were considered, reaching its maximum when the complete protein sequence was used (**Supplementary Fig. S16**). This indicates that mutation-associated decreases in predicted RNA-binding propensity are not always confined to a narrow window around the substituted residue. Some mutations induce broader changes in the predicted profile, consistent with local physicochemical perturbations propagating to nearby or distal regions of the sequence.

Based on this analysis, the default HERCULES mutation score was defined as the difference between the mean mutant and wild-type residue-level profiles across the complete sequence. This global summary provides a compact descriptor for mutation ranking and benchmarking, while the full residue-level profiles remain the primary output for interpreting where the predicted changes occur. The same definition can be applied to single substitutions, multiple substitutions and length-altering variants, for which a fixed local window may not capture all affected residues.

**Protein–RNA complex dataset**

On the protein–RNA complex dataset, we evaluated algorithm performance by analyzing the highest- and lowest-scoring residues for each protein. Specifically, for each method we selected the top and bottom 20% of residue scores on a per-protein basis and compared the resulting score distributions after normalizing all residue-level profiles to ensure direct comparability across algorithms. This analysis assesses the reliability of each method at the extremes of the score distribution, evaluating how consistently high- and low-confidence predictions correspond to true RNA-binding and non-binding residues, respectively.

**Exploratory analysis of PDB protein–RNA complexes and AlphaFold3-based contact augmentation**

We performed an exploratory analysis using experimentally resolved protein–RNA complexes from the Protein Data Bank (PDB), derived from previously curated protein–RNA complex datasets[3,4] (see the **Datasets** section in the **Methods**), to assess how HERCULES residue-level RNA-binding propensity profiles relate to atomic contact annotations derived from individual structural complexes. This analysis was not used for model training, parameter selection or threshold optimization.

The PDB-derived dataset differs substantially from the Pfam and InterPro RNA-binding region benchmarks used in the main residue-level validation. PDB entries often correspond to isolated domains, structured fragments or crystallizable regions rather than full-length proteins. Accordingly, proteins in the PDB benchmark are generally shorter than those in the Pfam benchmark, and the fraction of residues annotated as RNA-contacting is lower because only residues in direct contact with the RNA molecule present in the solved structure are labelled as positives. The dataset also shows biases in amino-acid composition, including enrichment of positively charged residues among RNA-contacting positions, consistent with the electrostatic nature of many protein–RNA interfaces. These dataset-level differences are summarized in **Supplementary Fig. S8**.

A key limitation of single-complex contact annotations is that they provide a sparse and context-specific view of RNA binding. In the curated PDB dataset, each entry reports the interaction between one protein chain and one RNA molecule. Residues that do not contact the RNA in that specific structure are therefore labelled as negatives, even though the same protein may use additional regions to bind different RNA ligands, RNA conformations or interaction contexts. This can introduce false-negative labels when evaluating methods designed to estimate general RNA-binding propensity rather than the contact map of one specific protein–RNA complex.

To explore this limitation, we generated additional protein–RNA complex models using AlphaFold3[5] for a subset of proteins from the PDB benchmark. AlphaFold3 was used only for exploratory contact augmentation and not as a source of definitive ground-truth labels. Each selected protein was modelled in complex with a panel of structured G-quadruplex-forming RNAs selected to increase the diversity of possible RNA-binding contexts[6,7]. The RNA panel included:

- gquad1: GGCUGGGGGAGGGGGCCGGGG
- G4A4: AAAAAAGGGGAAAAGGGGAAAAGGGGAAAAGGGGAAAAAA
- TERRA: UUAGGGUUAGGGUUAGGGUUAGGG
- VEGFA: GGAGGAGGGGGAGGAGGAAGA

AlphaFold3 predictions of protein–RNA complexes are not equivalent to experimentally determined structures and can be uncertain, particularly for flexible nucleic acids and non-canonical interfaces. We therefore retained these models only as exploratory structural evidence. For the predicted complexes considered here, the distributions of per-residue pLDDT values for the protein and RNA chains are shown in **Supplementary Fig. S9**. These values indicate moderate confidence for the retained models and were used to support their inclusion in the exploratory analysis.

For each protein–RNA pair, a single AlphaFold3 prediction run was performed, and all output structural models from that run were retained. Residue-level contact masks were generated by assigning a protein residue as RNA-contacting if any heavy atom of that residue was within 7 Å of any RNA atom in at least one retained model. Experimental PDB contact masks and AlphaFold3-derived contact masks were then combined to generate an augmented annotation for each protein. This augmented annotation was intended to increase the diversity of RNA interaction contexts represented for each protein, not to replace the experimentally determined PDB contacts.

Predictive performance was evaluated using a top-versus-bottom AUROC strategy, as previously used for residue-level propensity evaluation[2]. For each method and each protein, residues were ranked by predicted RNA-binding propensity, and AUROC was computed by comparing the highest-scoring and lowest-scoring 20% of residues. Performance was first evaluated using only experimental PDB contact annotations and then using the augmented annotations incorporating AlphaFold3-derived G-quadruplex contacts. HERCULES, DRNApred[8] and HydRA[1] were included in this comparison. When evaluated only on experimental PDB contacts, all methods showed modest performance, consistent with the sparsity and context dependence of single-complex contact maps. After augmentation with AlphaFold3-derived contacts, HERCULES showed a modest increase in median AUROC, whereas DRNApred and HydRA generally decreased (**Supplementary Fig. S10**). This result suggests that some high-scoring HERCULES regions may be more consistent with expanded RNA-binding contexts than with a single structural contact map alone. However, because the augmented contacts are predicted rather than experimentally determined, this analysis should be interpreted as exploratory and hypothesis-generating.

Finally, we assessed sequence overlap between the PDB benchmark and the training sets of PDB-trained residue-level predictors. NCBRPred[9] showed near-complete sequence redundancy with the test proteins, whereas DRNApred showed lower overlap (**Supplementary Fig. S11**). For this reason, NCBRPred was not included in the primary structural benchmark. Overall, the PDB/AlphaFold3 analysis is reported as a complementary assessment of contact-label incompleteness and RNA-context dependence, rather than as a principal validation of HERCULES residue-level performance.

##### Prediction of experimental ΔΔG from HERCULES profiles

To investigate whether HERCULES residue-level profiles encode quantitative information about mutation-induced changes in protein–RNA binding affinity, we trained a separate Gradient Boosting regression model to predict experimental binding free-energy changes (ΔΔG). This analysis was performed as an auxiliary evaluation of the information contained in HERCULES profiles and is not part of the default HERCULES output.

The regression model used the complete wild-type and mutant HERCULES residue-level profiles as input features. Because protein sequences have variable length, profiles were resampled to a fixed-length representation before concatenating wild-type and mutant profiles into a single feature vector for each mutation. The model was trained on the PRA-MutPred dataset[10], using protein-level splits to prevent information leakage between mutations occurring in the same protein. Hyperparameters were optimized using the training set, and performance was evaluated on a held-out test set. Model performance was quantified using Pearson and Spearman correlation coefficients between predicted and experimentally measured ΔΔG values (**Supplementary Fig. S18**). As an additional application, the trained regression model was applied without retraining to the independent UniProt mutation dataset used for the HERCULES mutation-effect analysis (**Supplementary Fig. S19**). This dataset contains experimentally validated RNA-binding-disrupting mutations from diverse proteins, including both single and multiple amino-acid substitutions, but does not provide experimental ΔΔG values. We therefore used this analysis only to examine whether mutations annotated as impairing RNA binding were assigned positive predicted ΔΔG values, consistent with reduced binding affinity.

### Supplementary Tables

**Supplementary Table S1.** Global RNA-binding propensity scores computed by HERCULES for all reviewed human UniProt entries. The table includes annotation indicating whether each protein was included in the training or independent test sets used for binary RBP classification.

**Supplementary Table S2.** Dataset of human proteins containing Pfam-annotated RNA-binding domains (RBDs), classified as RNABinding, NonClassical, or UnknownPutative.

**Supplementary Table S3.** Dataset of human proteins containing RNA-binding domains (RBDs) from the InterPro database.

**Supplementary Table S4.** Dataset of experimentally validated mutations reported to impair RNA-binding activity, retrieved from UniProt and manually curated.

**Supplementary Table S5.** Mapping of feature names and categorization in feature families.

**Supplementary Table S6.** Dataset of proteins extracted from experimentally determined protein–RNA complexes deposited in the Protein Data Bank (PDB).

**Supplementary Table S7.** Raw protein-array measurements for the three repeat-RNA probes (CGG, CAG and CAACAG); each lists all mapped proteins on the array. For every protein the table reports the gene symbol, UniProt accession and protein name, together with the three replicate binding signals (Signal_rep1–3) and their mean (Mean_signal).

**Supplementary Table S8.** 18 HERCULES candidate RBPs positive in protein-microarray experiments.

### Supplementary Figures

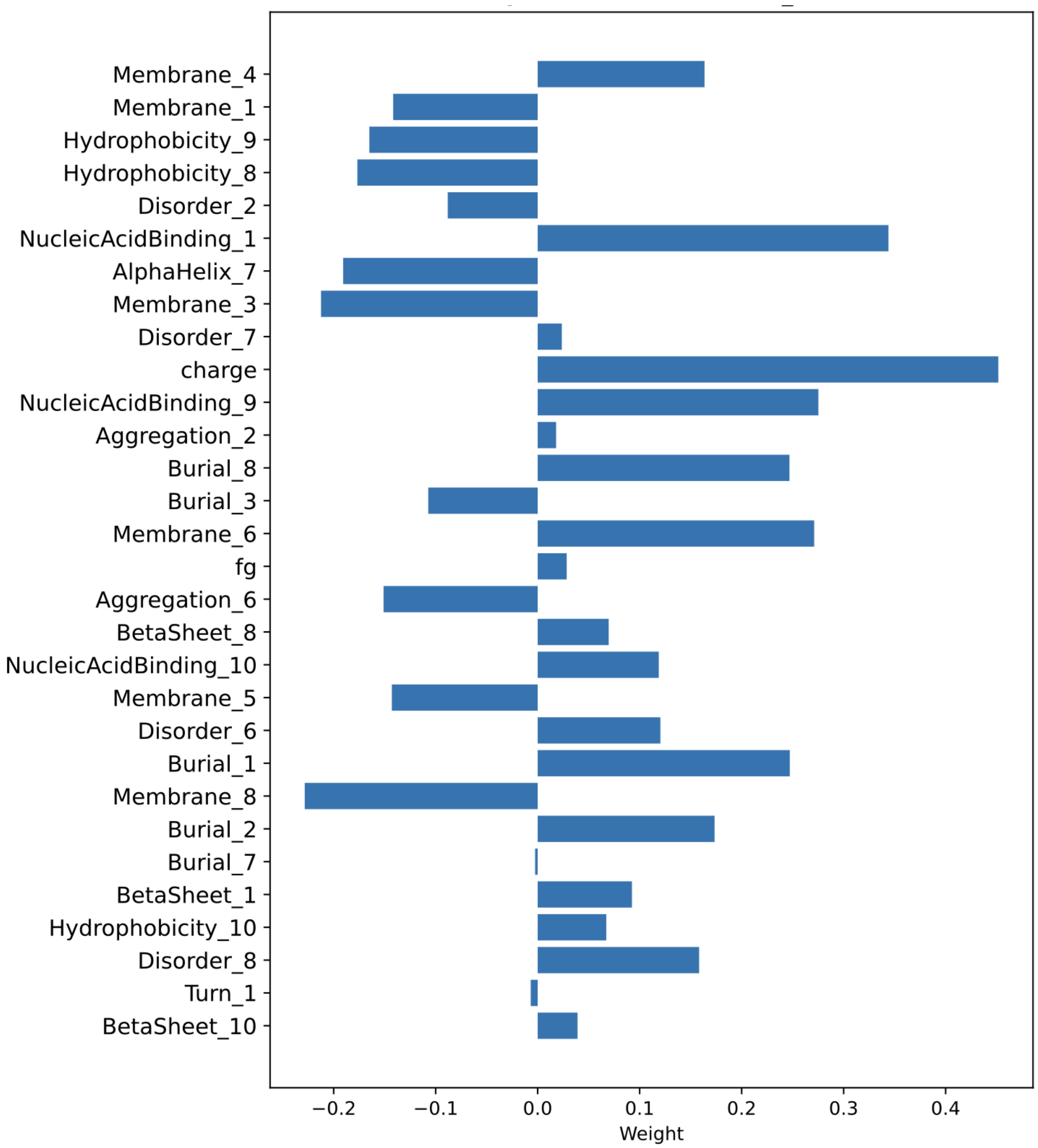


**Supplementary Figure S1.** Bar plot showing the weights of the selected physicochemical features, illustrating their relative contributions to the local physicochemical information used to quantify RNA-binding propensity. The features were selected from a pool of physicochemical scales reported in **Supplementary Table S5**, which contains a collection of 10 predictors for each category.

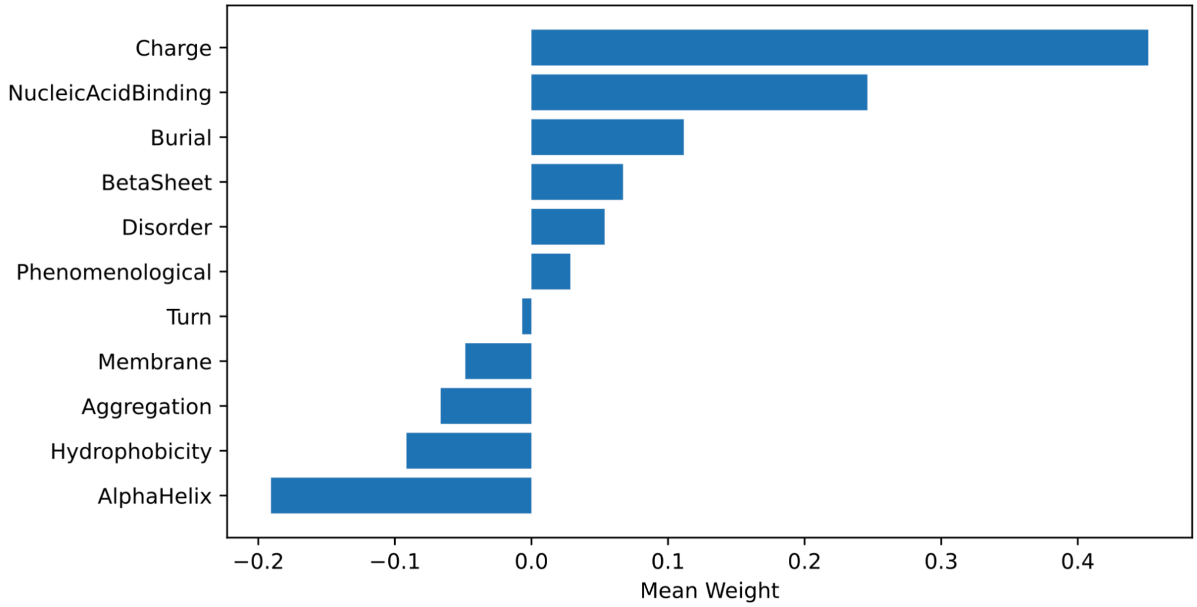


**Supplementary Figure S2.** Contribution of the differences in physicochemical properties that are used to score the model, in particular the mutation effect.

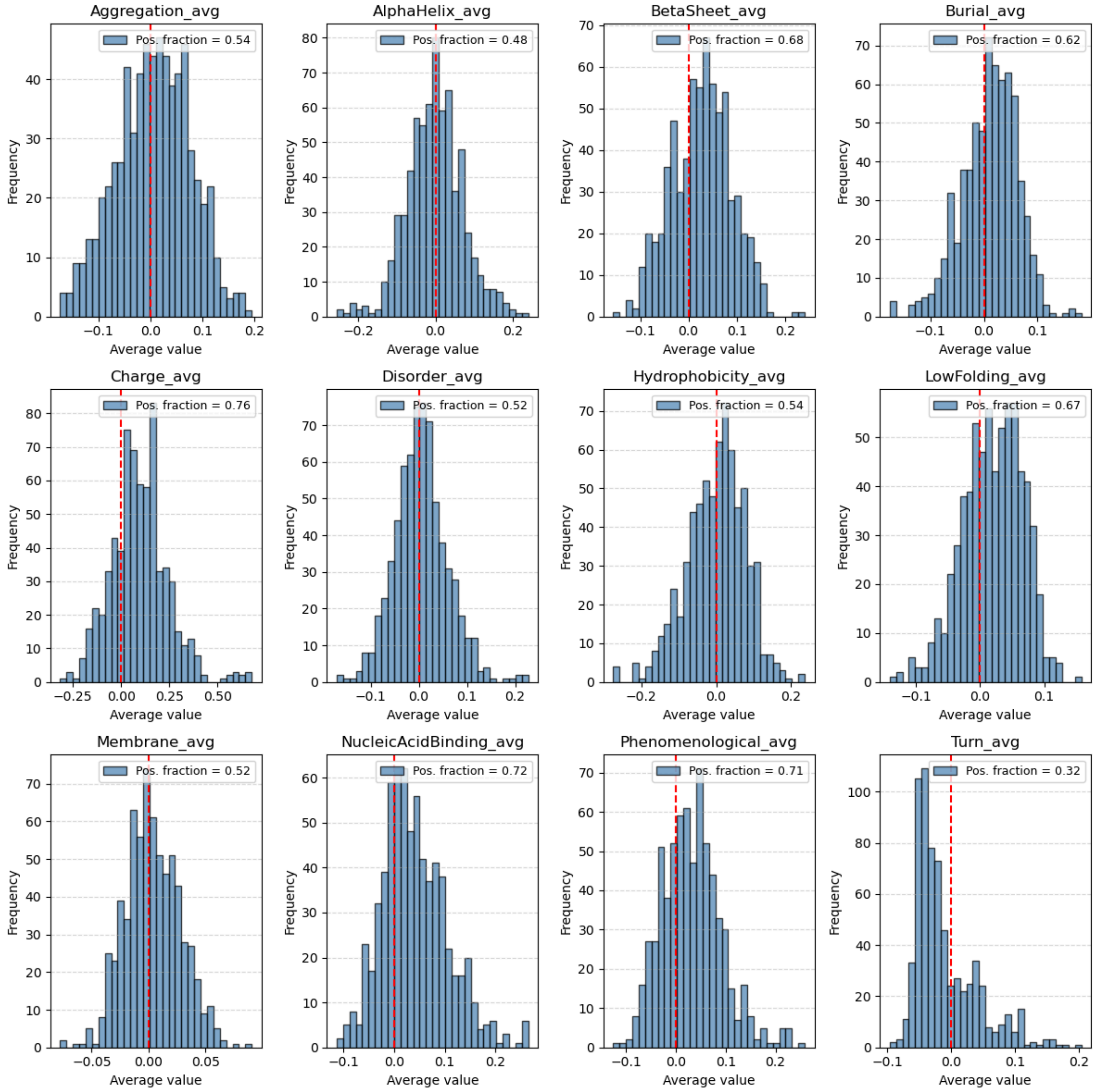


**Supplementary Figure S3.** Histograms of the differences in physicochemical properties between the wild-type (WT) and mutated proteins, highlighting the fraction of positive differences. No single averaged property alone is able to reproduce the same performance in recognizing deleterious mutations as the combined linear model that integrates all properties.

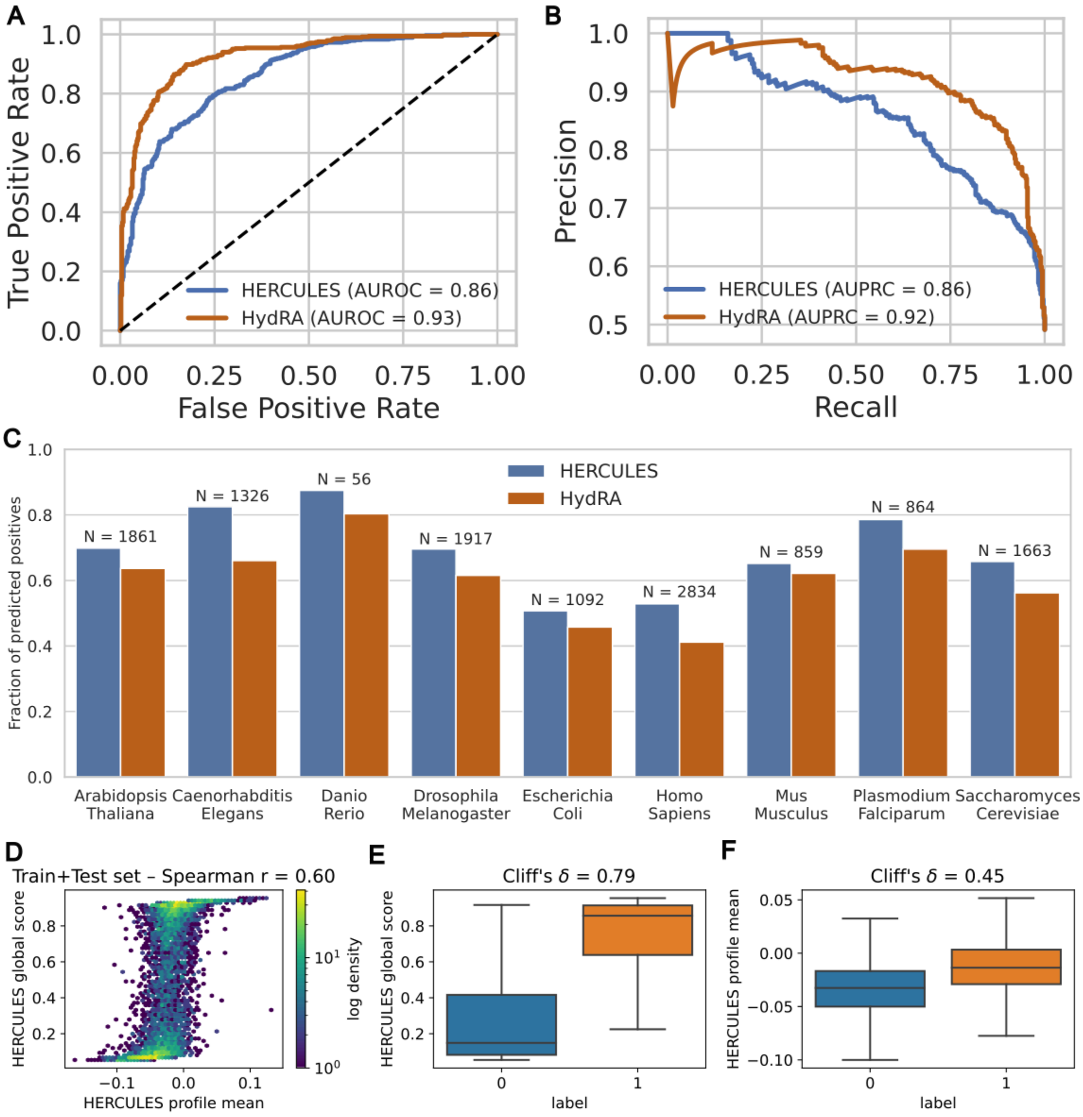


**Supplementary Figure S4.** Performance of the HERCULES global score in predicting RNA-binding proteins. **(A)** Receiver operating characteristic (ROC) curves comparing HERCULES and HydRA on an independent human test set, filtered with <30% sequence identity with the HERCULES training set (see **Methods**). The dashed line indicates random performance. **(B)** Precision–recall curves for the same dataset. **(C)** Fraction of predicted RBPs across nine organisms using RBPs curated from RBP2GO, filtered with <30% sequence identity with the HERCULES training set (see **Methods**). Bars show the proportion of proteins classified as RNA-binding by binarized HERCULES and HydRA scores; the number of proteins analyzed in each organism is indicated above each bar. **(D)** Density-colored scatter plot showing the relationship between the HERCULES global score and the mean of the residue-level RNA-binding propensity profile on the train–test set (Spearman ρ = 0.59). **(E)** Box plots illustrating separation of RBPs (label = 1) and non-RBPs (label = 0) by the HERCULES global score, with a Cliff's δ effect size indicated. **(F)** Box plots showing separation of RBPs and non-RBPs by the mean of the residue-level HERCULES profile.

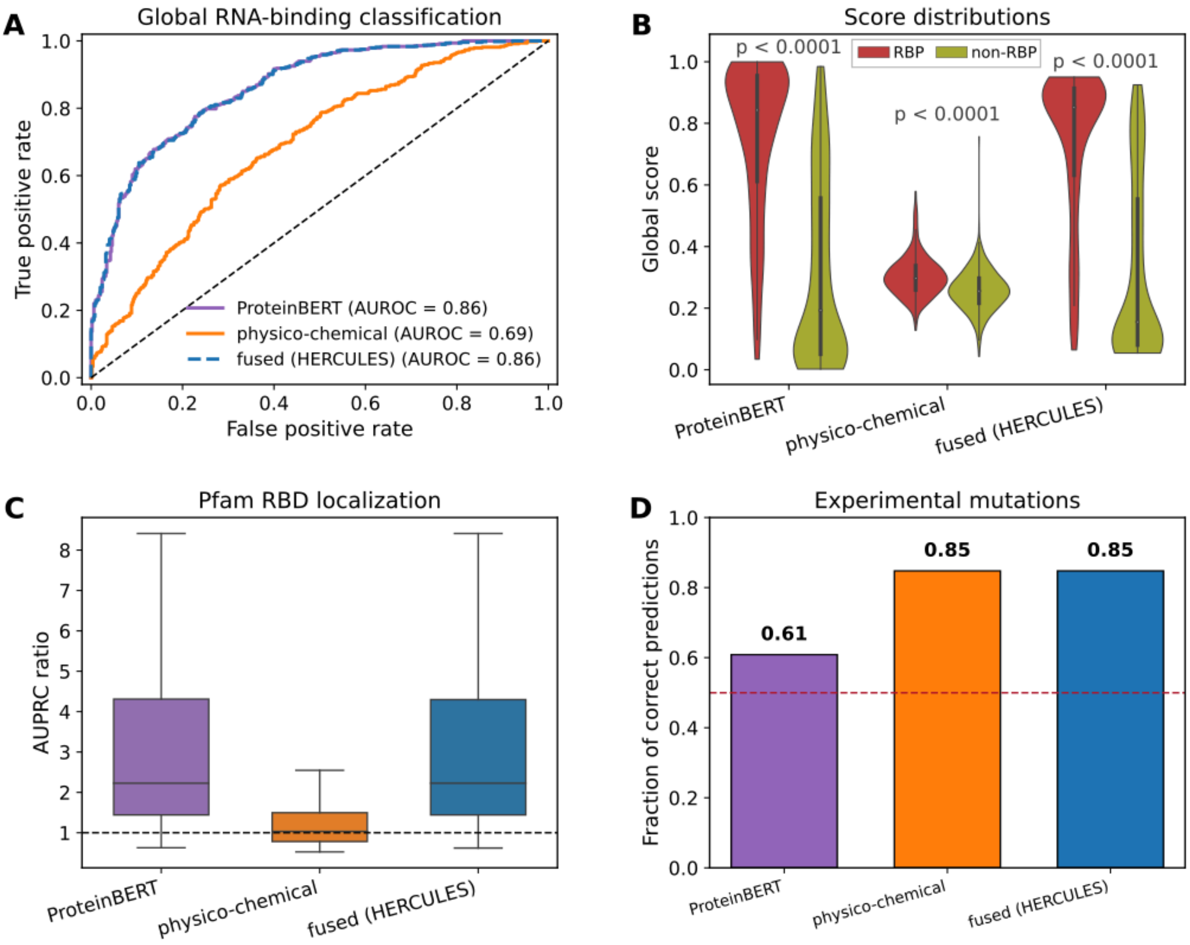


**Supplementary Figure S5. Ablation of HERCULES model branches across global, residue-level and mutation prediction tasks. (A)** Receiver operating characteristic (ROC) curves for global RNA-binding protein (RBP) classification on an independent test set of 979 proteins (<30% sequence identity to the training set). Area under the ROC curve (AUROC) is shown in the legend. The dashed diagonal indicates random performance. **(B)** Violin plots of global scores stratified by RBP (red) and non-RBP (olive) labels for each branch. Two-sided Mann–Whitney U tests indicate significant separation between RBP and non-RBP distributions for all three branches ($P < 0.0001$) **(C)** Box plots of the per-protein area under the precision–recall curve (AUPRC) ratio for Pfam RNA-binding domain (RBD) localization on a retained validation set of 604 proteins (<30% identity to training). The dashed horizontal line at 1 denotes random performance. **(D)** Bar plot showing the fraction of correctly predicted experimentally annotated mutations from the independent test set (n = 92, 28 proteins). The dashed red line at 0.5 indicates random expectation.

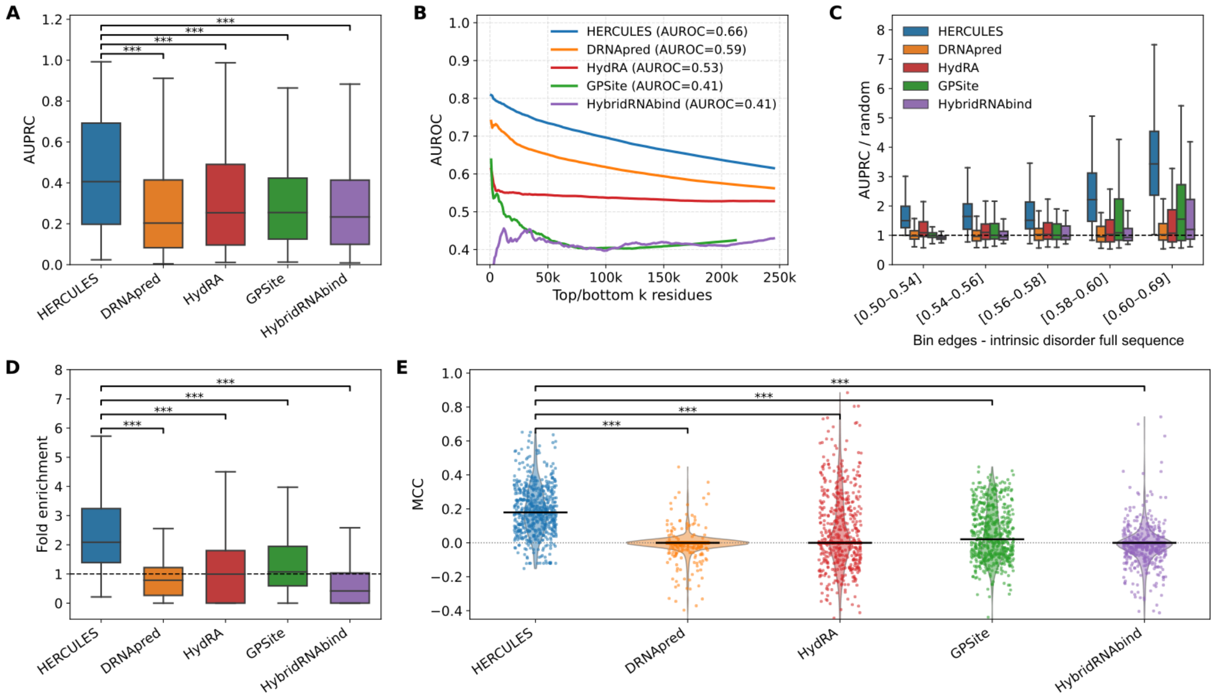


**Supplementary Figure S6. Additional evaluation of residue-level RNA-binding predictions on the Pfam-annotated benchmark. (A)** Box plots of raw per-protein area under the precision–recall curve (AUPRC) for HERCULES, DRNApred, HydRA, GPSite and HybridRNAbind on the Pfam RBD validation set used in Figure 2 (n = 604 proteins). **(B)** Area under the receiver operating characteristic curve (AUROC) computed as a function of the number of top- and bottom-ranked residues (k); legend values report the AUROC on the full protein set. **(C)** Box plots of the AUPRC ratio to random stratified by binning the values of the mean intrinsic disorder over the full protein sequence (DisProt scale). **(D)** Box plots showing the fold enrichment of RNA-binding residues within the top 10% highest-scoring positions; the dashed line indicates no enrichment (fold = 1). **(E)** Raincloud plots (violin density, per-protein points and median bar) of Matthews correlation coefficient (MCC) using method-appropriate binary classification thresholds: top 10% of scores for HERCULES and GPSite; author-defined native binary profiles for DRNApred and HybridRNAbind; and HydRA occlusion significance peaks ($p \leq 0.05$). Brackets in **(A)**, **(D)** and **(E)** denote Bonferroni-adjusted Wilcoxon signed-rank tests for paired per-protein comparisons of HERCULES versus each algorithm (***padj < 0.001).

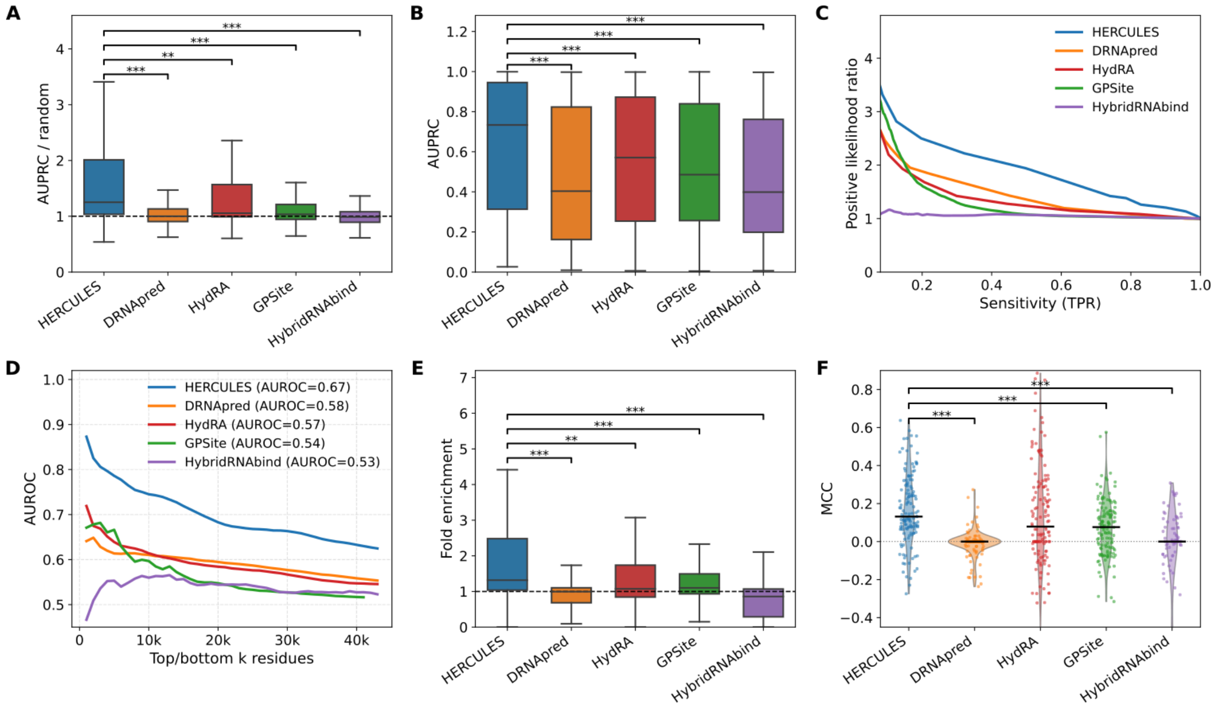


**Supplementary Figure S7.** Additional evaluation of residue-level RNA-binding predictions on the InterPro non-Pfam benchmark. (A) Box plots showing the ratio of per-protein area under the precision–recall curve (AUPRC) to a random baseline (AUPRC/random) for HERCULES, DRNApred, HydRA, GPSite and HybridRNAbind on the independent InterPro non-Pfam RBD validation set filtered to <30% sequence identity to the HERCULES training set (n = 140 proteins); the dashed line indicates equivalence with random expectation (ratio = 1). (B) Box plots of raw per-protein AUPRC for the same methods and dataset. (C) Positive likelihood ratio (PLR) as a function of sensitivity (true positive rate), computed from residue-level scores and labels pooled across all benchmark proteins. (D) Area under the receiver operating characteristic curve (AUROC) computed as a function of the number of top- and bottom-ranked residues (k) for each method; legend values report full-sequence AUROC. (E) Box plots showing the fold enrichment of RNA-binding residues within the top 10% highest-scoring positions (dashed line, no enrichment). (F) Raincloud plots (violin density, per-protein points and median bar) of Matthews correlation coefficient (MCC) using method-appropriate binary classification thresholds: top 10% of scores for HERCULES and GPSite; author-defined native binary profiles for DRNApred (binary_RNA) and HybridRNAbind (RNA_Binary); and HydRA occlusion significance peaks ($p \leq 0.05$). Brackets in (A), (B), (E) and (F) denote Bonferroni-adjusted Wilcoxon signed-rank tests for paired per-protein comparisons of HERCULES versus each comparator (*padj < 0.05, **padj < 0.01, ***padj < 0.001).

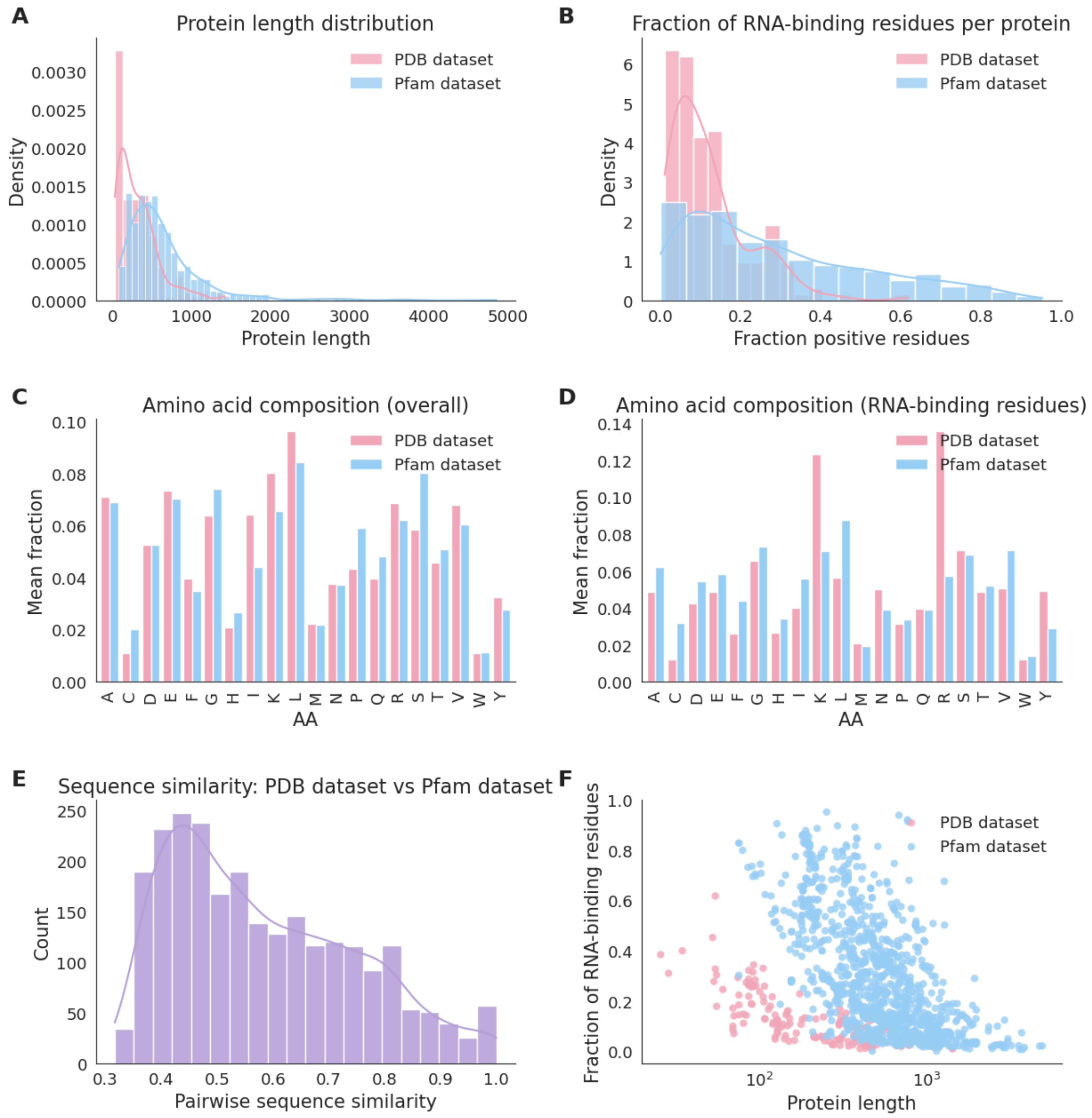


**Supplementary Figure S8.** Comparison between Pfam and PDB datasets. **(A)** Distribution of protein lengths in the PDB and Pfam datasets. **(B)** Distribution of the fraction of residues annotated as RNA-binding per protein. **(C)** Mean amino acid composition computed over full protein sequences. **(D)** Mean amino acid composition restricted to RNA-binding residues. **(E)** Distribution of pairwise sequence similarity scores computed between randomly sampled proteins from the two datasets. **(F)** Relationship between fraction of RNA-binding residues and protein length, shown for individual proteins.

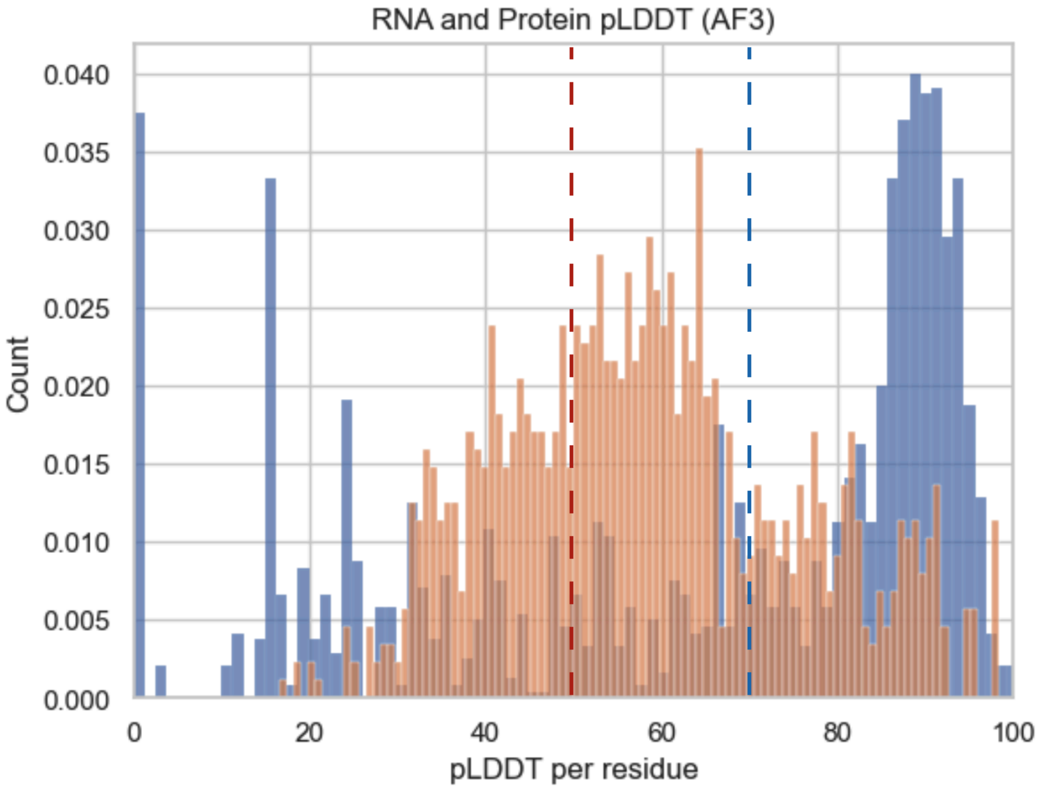


**Supplementary Figure S9.** Histogram of per-residue pLDDT scores for all protein–RNA complexes predicted by AF3 in association with G-quadruplex RNAs. The average pLDDT is 51 for the RNA components and 72 for the protein components. These values indicate that the selected predicted complexes exhibit overall moderate to good confidence, supporting their structural reliability for downstream analyses.

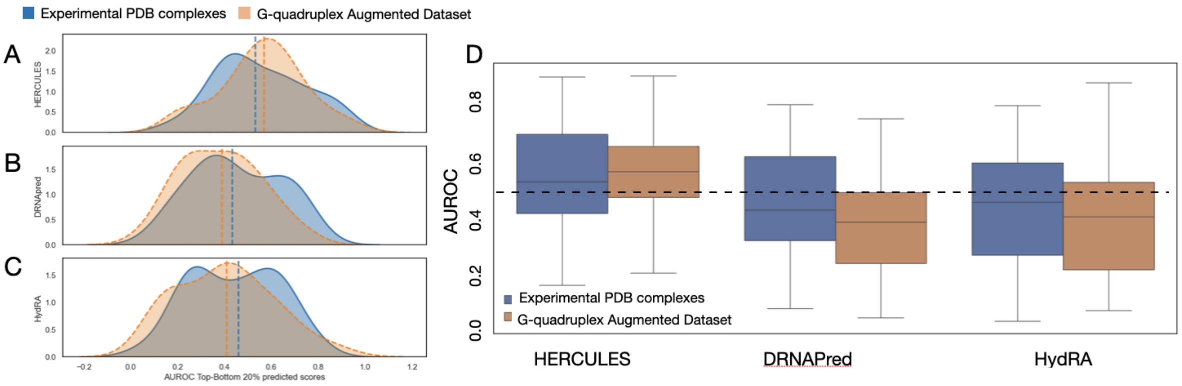


**Supplementary Figure S10. Exploratory evaluation of residue-level predictions on experimental and AlphaFold3-augmented protein–RNA contact annotations.** (A–C) Distributions of per-protein top-versus-bottom AUROC values for HERCULES (A), DRNApred (B) and HydRA (C). For each protein and method, AUROC was computed by comparing the top and bottom 20% of residues ranked by predicted RNA-binding propensity. Solid curves show performance using contact annotations from experimentally resolved PDB protein–RNA complexes, whereas dashed curves show performance after augmenting the same proteins with additional RNA-contact annotations derived from AlphaFold3-predicted complexes with multiple G-quadruplex RNAs. Vertical lines indicate the median of each distribution. (D) Box plot summarizing the AUROC distributions shown in panels A–C. HERCULES shows a modest increase in median AUROC after contact augmentation, whereas DRNApred and HydRA decrease, consistent with the interpretation that HERCULES high-scoring regions are more compatible with expanded RNA-binding contexts than with a single experimental contact map alone.

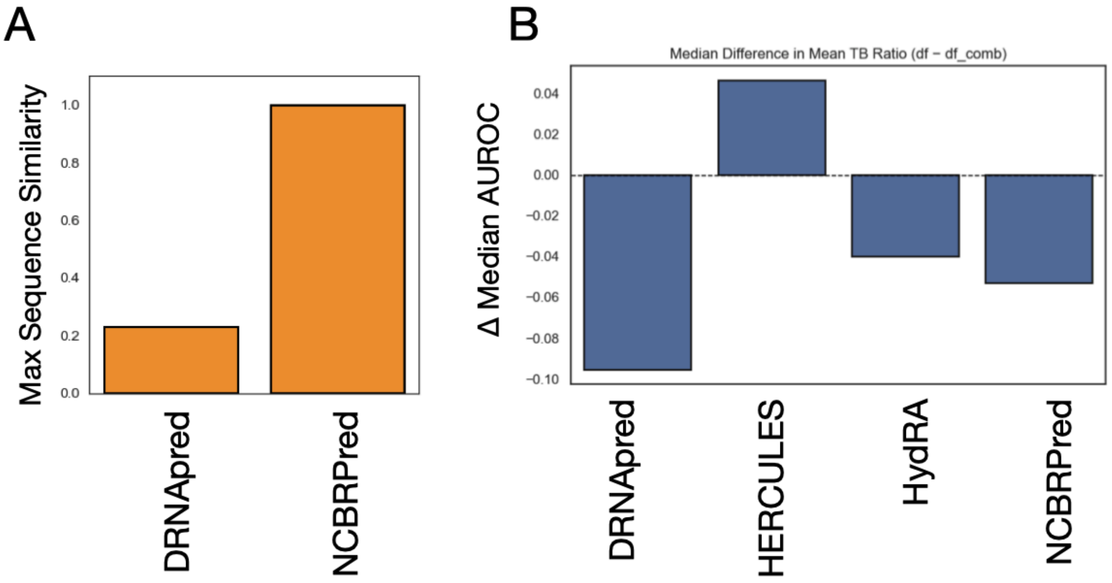


**Supplementary Figure S11. Sequence-redundancy analysis for PDB-trained residue-level predictors and comparison of experimental versus AlphaFold3-augmented contact annotations. (A)** Sequence similarity between the PDB benchmark proteins and the training datasets of DRNApred and NCBRPred. Because the NCBRPred training set shows near-complete sequence redundancy with the selected PDB test proteins, NCBRPred was excluded from the primary structural-complex benchmark. **(B)** Difference between the median scores of the top 20% and bottom 20% of residues for experimental PDB contact annotations and AlphaFold3-augmented annotations enriched for G-quadruplex RNA interactions.

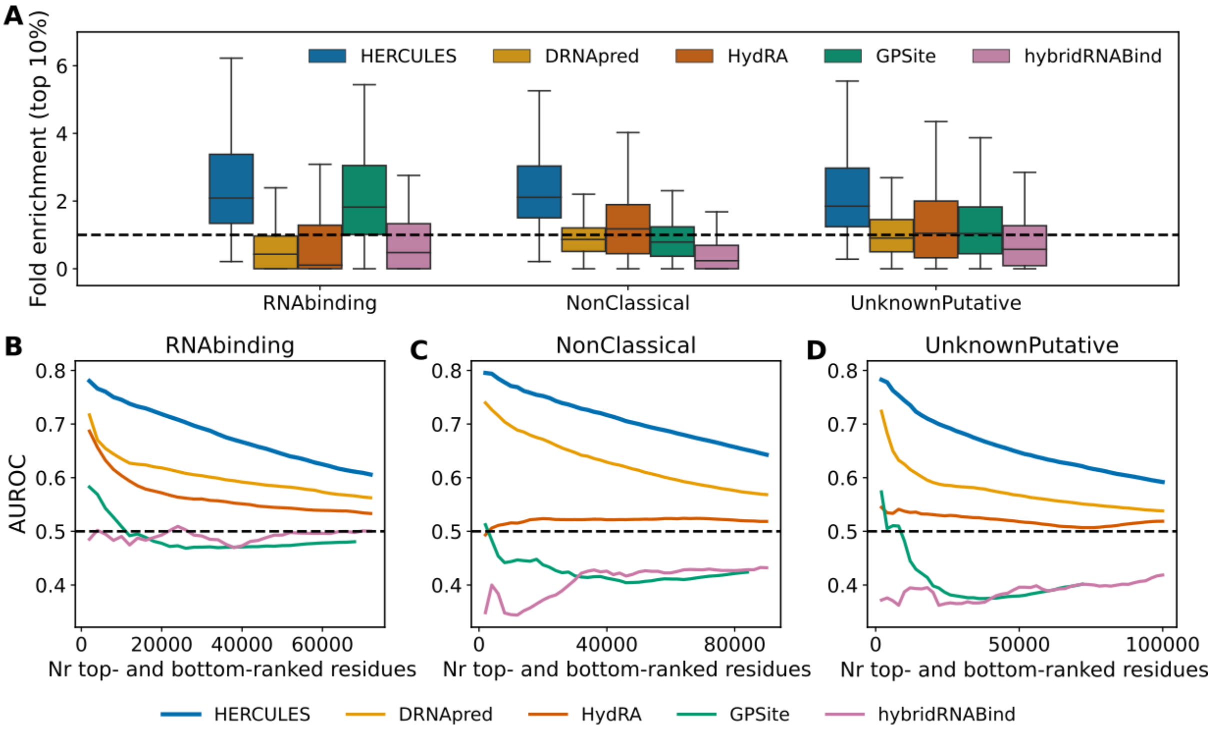


**Supplementary Figure S12.** Additional evaluation of predictions of canonical and non-canonical RBDs. **(A)** Box plots of fold enrichment in the top 10% highest-scoring residues for proteins grouped by RNA-binding domain class (RNABinding, NonClassical, UnknownPutative), corresponding to the dataset analyzed in Figure 3. **(B-D)** AUROC computed as a function of the number of top- and bottom-ranked residues (*k*) for each RNA-binding domain class. Each panel corresponds to one domain class and reports the performance of the five algorithms evaluated in panel A.

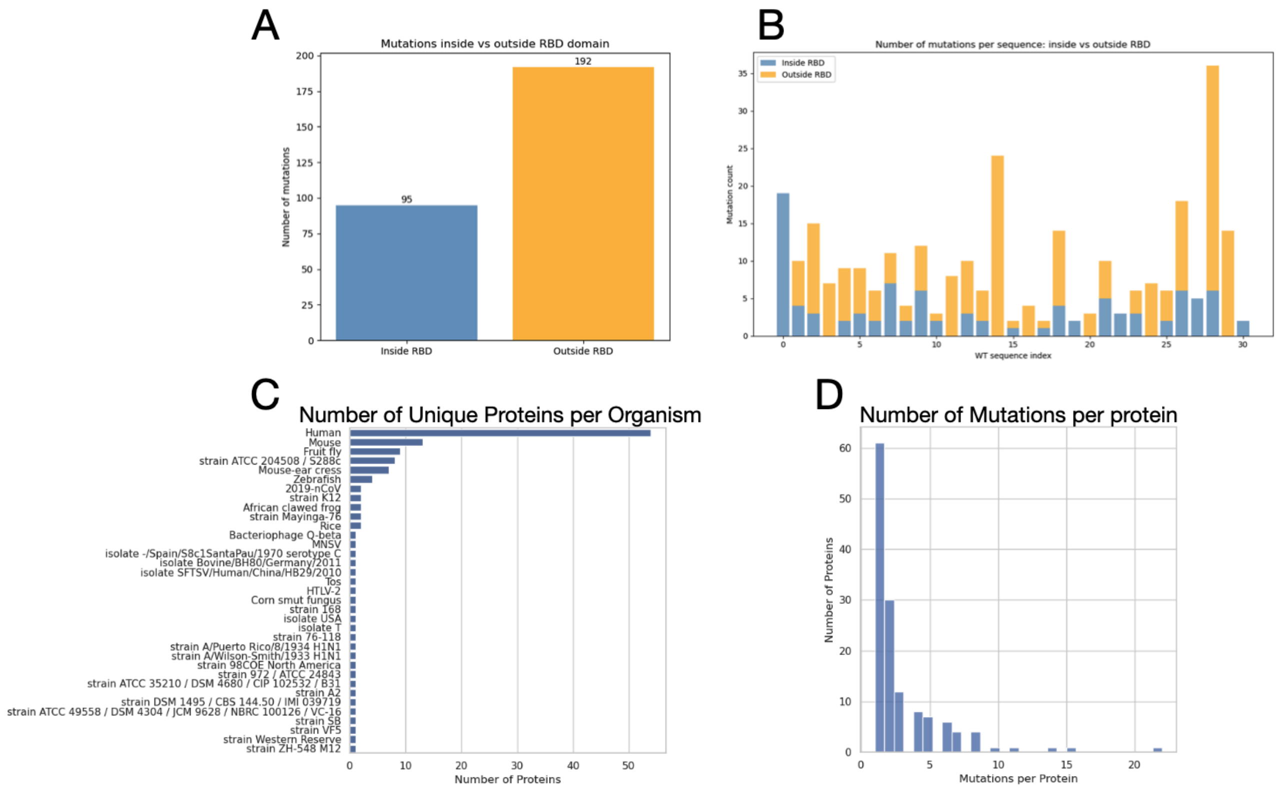


**Supplementary Figure S13.** Summary statistics of RNA-binding-disrupting mutations. **(A)** Bar plot showing the number of negative mutations occurring within and outside the RNA-binding domain (RBD) for human proteins. **(B)** Stacked bar plot showing the number of mutations inside (blue) or outside (yellow) the RBD for each human protein in our curated mutation dataset. **(C)** Bar plot showing the number of proteins per organism in the curated mutation dataset. **(D)** Histogram of the number of mutations per protein.

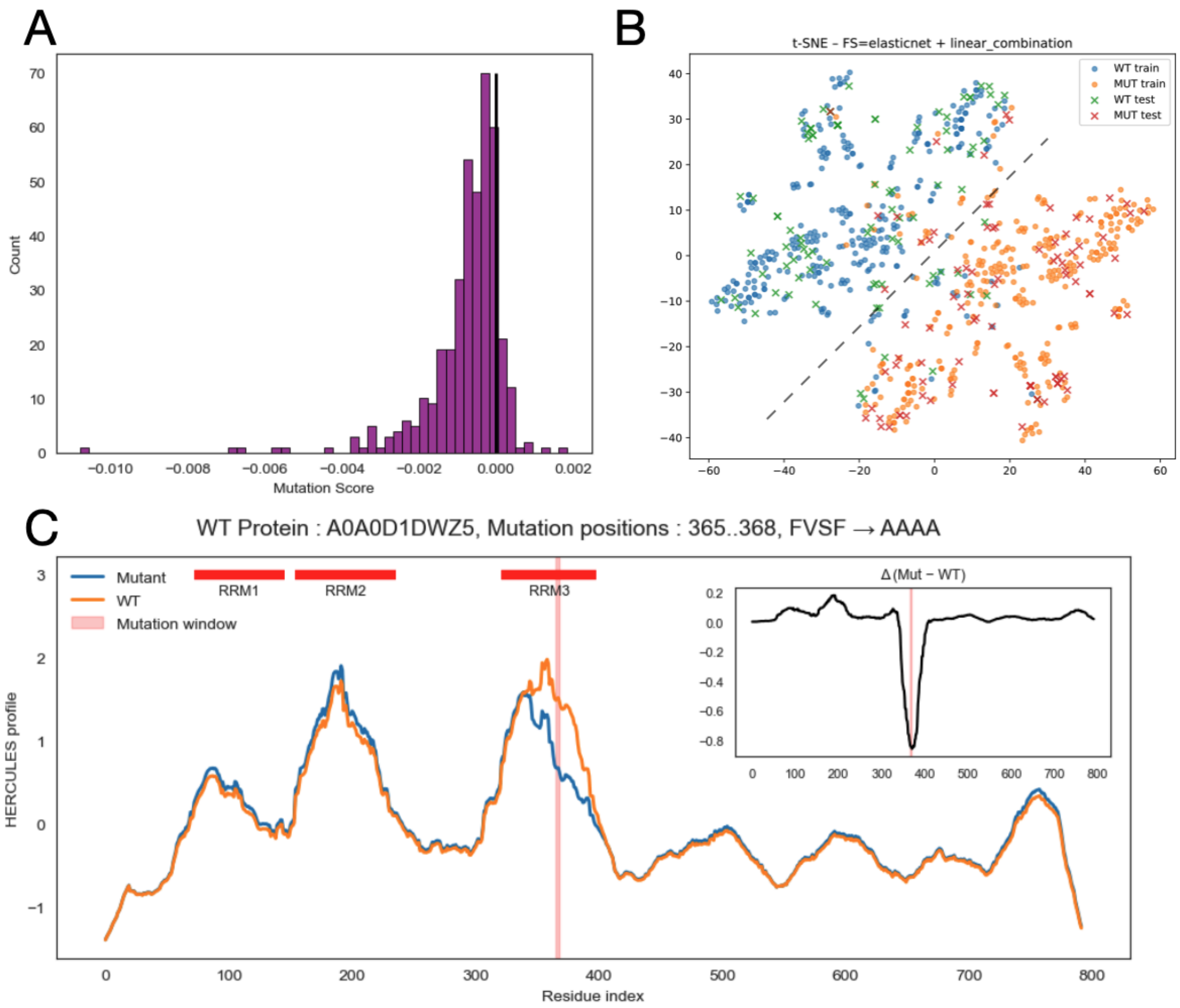


**Supplementary Figure S14. (A)** Histogram of mutation scores, defined as the difference between the means of the mutant and wild-type (WT) RNA-binding propensity profiles (mutant-WT), computed for all variants in the held-out test set of the curated mutation dataset. The fraction of mutations predicted to cause loss of RNA binding (negative scores) is 0.87. **(B)** *t-SNE* representation of the score difference between mutated and wild-type (WT) sequences. Dots indicate training samples, while crosses denote test samples. **(C)** HERCULES residue-level RNA-binding propensity profiles for the WT (orange) and mutant (blue) sequences around the mutation region. The example protein (UniProt ID A0A0D1DWZ5) carries a four–amino-acid substitution (positions 365–368, FVSF → AAAA). The inset shows the profile difference (mutant − WT), highlighting a localized decrease in RNA-binding propensity at the mutation sites.

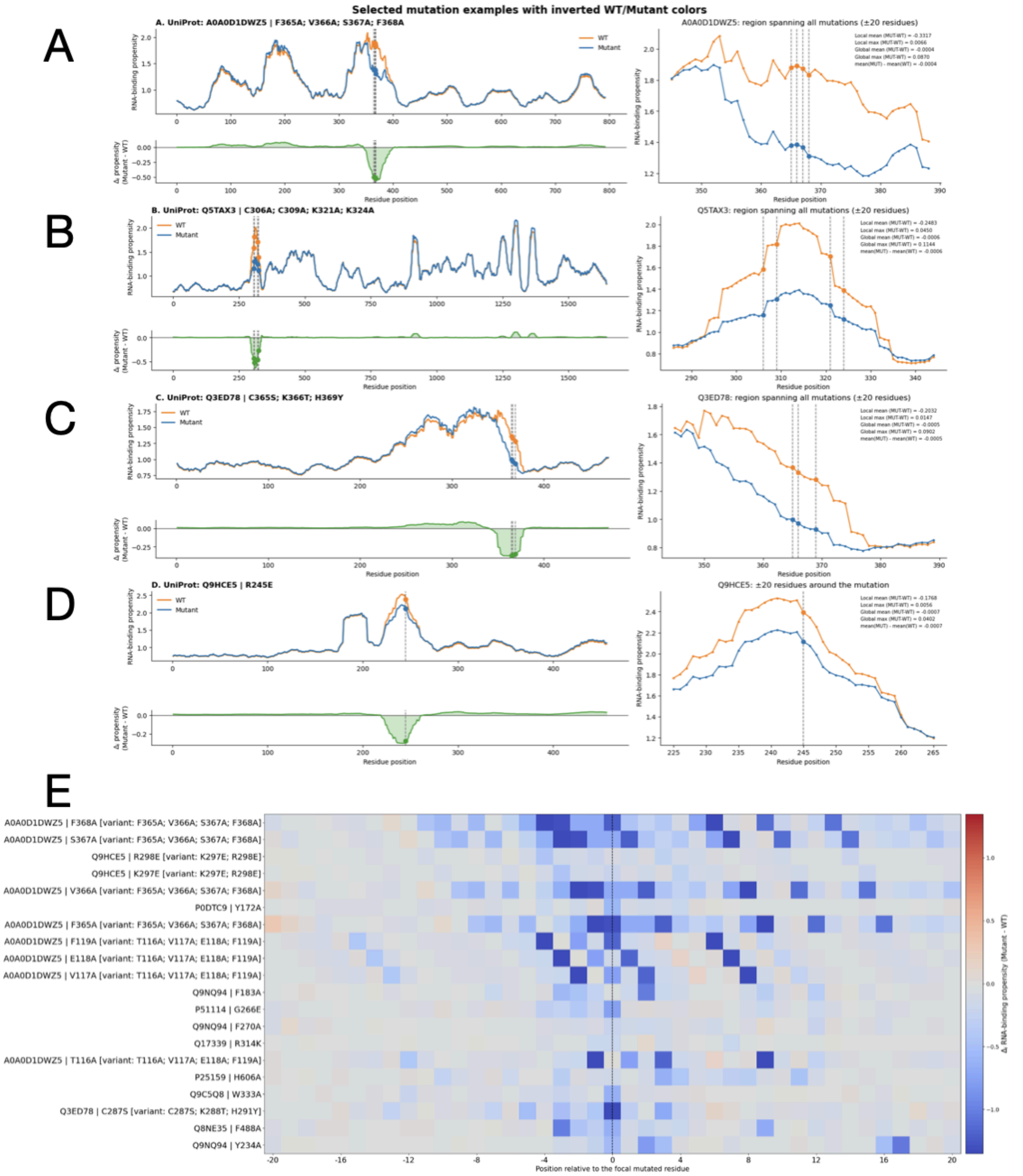


**Supplementary Figure S15**. **Predicted residue-level RNA-binding propensity changes for experimentally validated RNA-binding–abolishing mutations**. Panels (**A–D**) show four representative variants selected for consistent predicted loss of binding propensity. For each variant, left: full-length WT and mutant profiles (25-aa smoothed; dashed lines, mutation sites); bottom left: Mutant − WT difference; right: local zoom (±20 aa) with summary statistics. Panel **E**: mutation-centered heatmap of WT − Mutant propensity differences for 20 additional variants with the largest predicted global effects (unsmoothed profiles). All mutations shown experimentally abolish RNA binding.

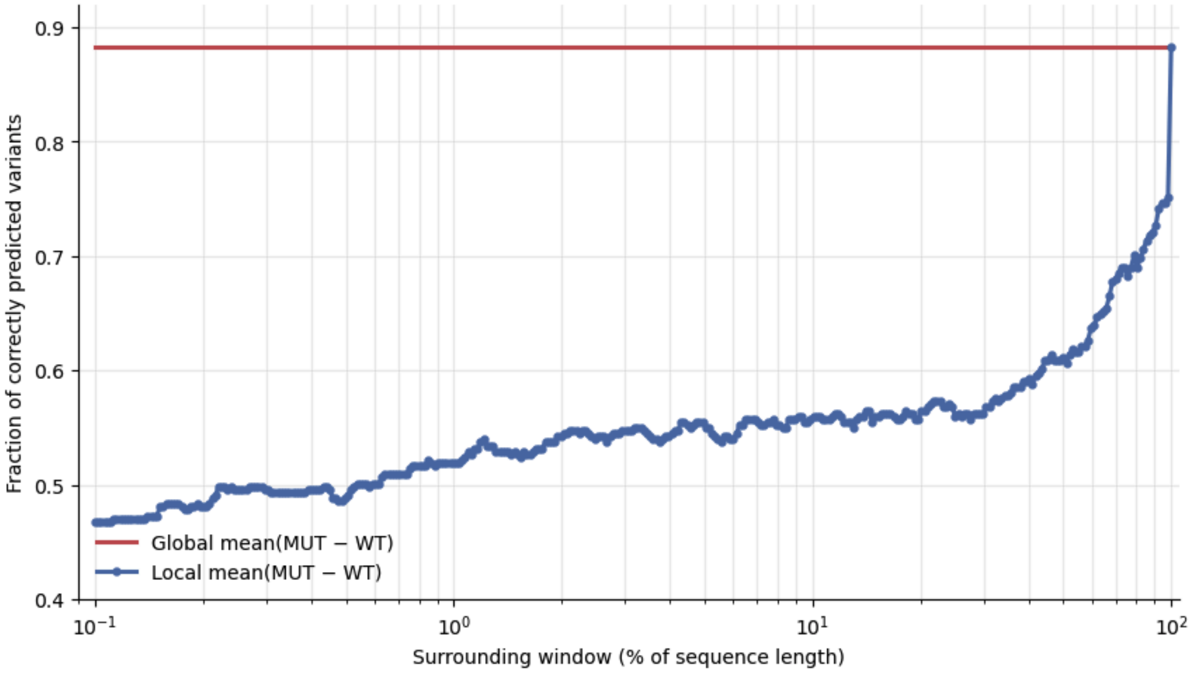


**Supplementary Figure S16**. **Comparison of global and local profile effects.** The fraction of deleterious (negative) mutations correctly identified from profile differences is plotted as a function of the sequence window surrounding the mutation site. The x-axis represents the distance from the mutation position, expressed as a percentage of the total protein length, while the y-axis reports the fraction of correctly predicted negative mutations.

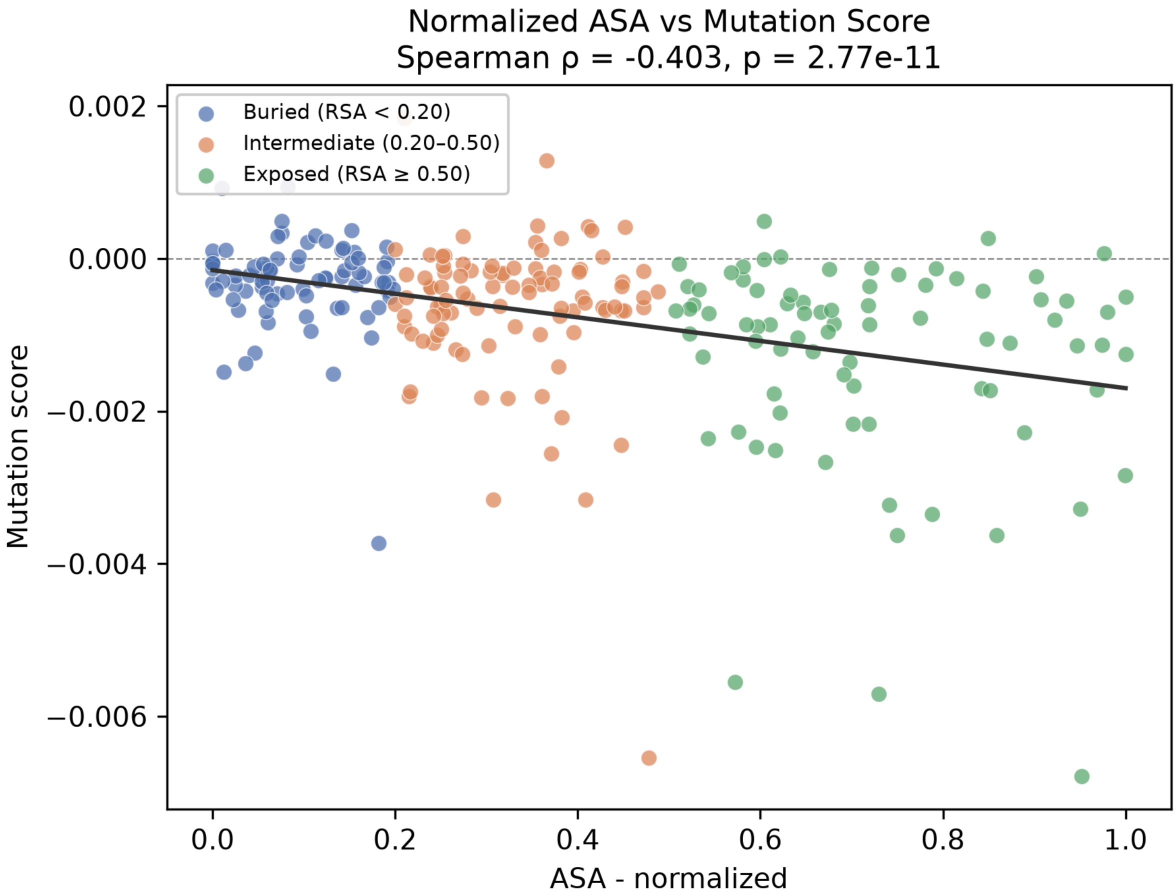


**Supplementary Figure S17. Influence of solvent accessibility on HERCULES mutation scores.** Relationship between HERCULES mutation score and mean relative solvent accessibility (RSA) at the substituted residues. RSA was computed as the solvent-accessible surface area (ASA) normalized by the maximum accessible surface area expected for each amino-acid type. Mutations are grouped as buried (RSA < 0.20), intermediate (0.20–0.50) or exposed (RSA ≥ 0.50). Higher RSA values are associated with more negative HERCULES mutation scores, indicating larger predicted decreases in RNA-binding propensity for substitutions affecting more solvent-exposed residues. This trend is consistent with the expected surface accessibility of many RNA-binding interfaces.

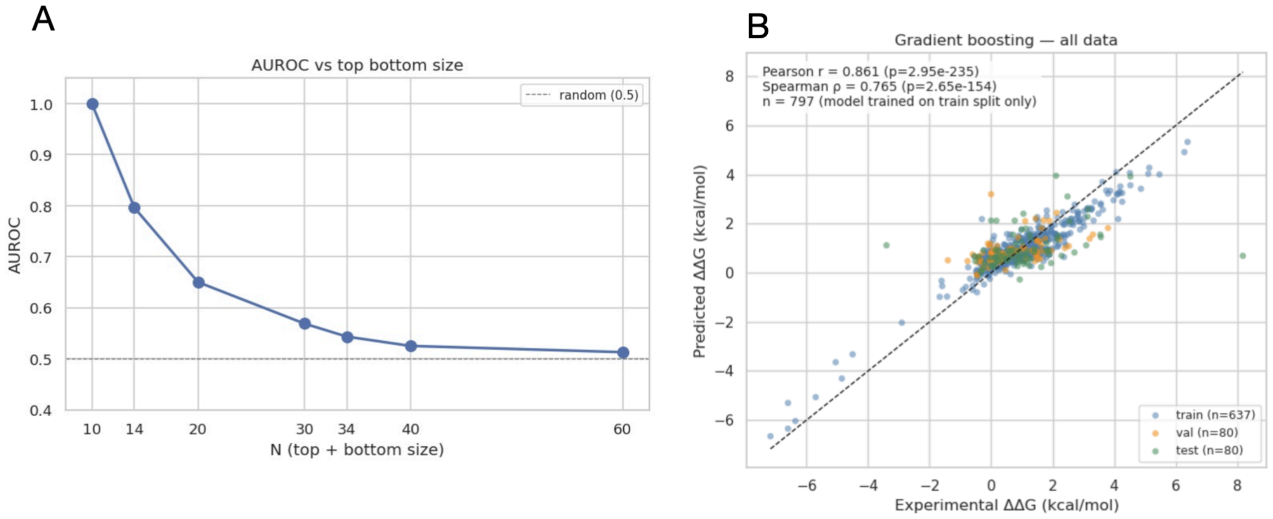


**Supplementary Figure S18. Quantitative prediction of RNA-binding affinity changes from HERCULES residue-level profiles.** (**A**) Top–bottom analysis of the HERCULES Δ-profile mean against experimental ΔΔG values from PRA-MutPred. Mutations were ranked according to their experimental ΔΔG (kcal/mol), and the top and bottom *N* mutations were selected. For each value of *N*, the AUROC was computed between the HERCULES Δ-profile mean binarized on the top and bottom N values and the experimental ΔΔG scores. The x-axis represents the number of top and bottom mutations included in the analysis, while the y-axis shows the AUROC. The analysis includes both stabilizing (negative ΔΔG) and destabilizing (positive ΔΔG) mutations. (**B**) Scatter plot comparing experimentally measured ΔΔG values with ΔΔG values predicted by a Gradient Boosting regression model trained using the wild-type and mutant HERCULES residue-level propensity profiles as input features. Each point represents a single mutation. The correlation between predicted and experimental ΔΔG indicates that quantitative information about mutation-induced changes in RNA-binding affinity is encoded within the HERCULES profiles, even though the default HERCULES mutation score is designed to detect disruption of RNA-binding capability rather than to directly estimate ΔΔG.

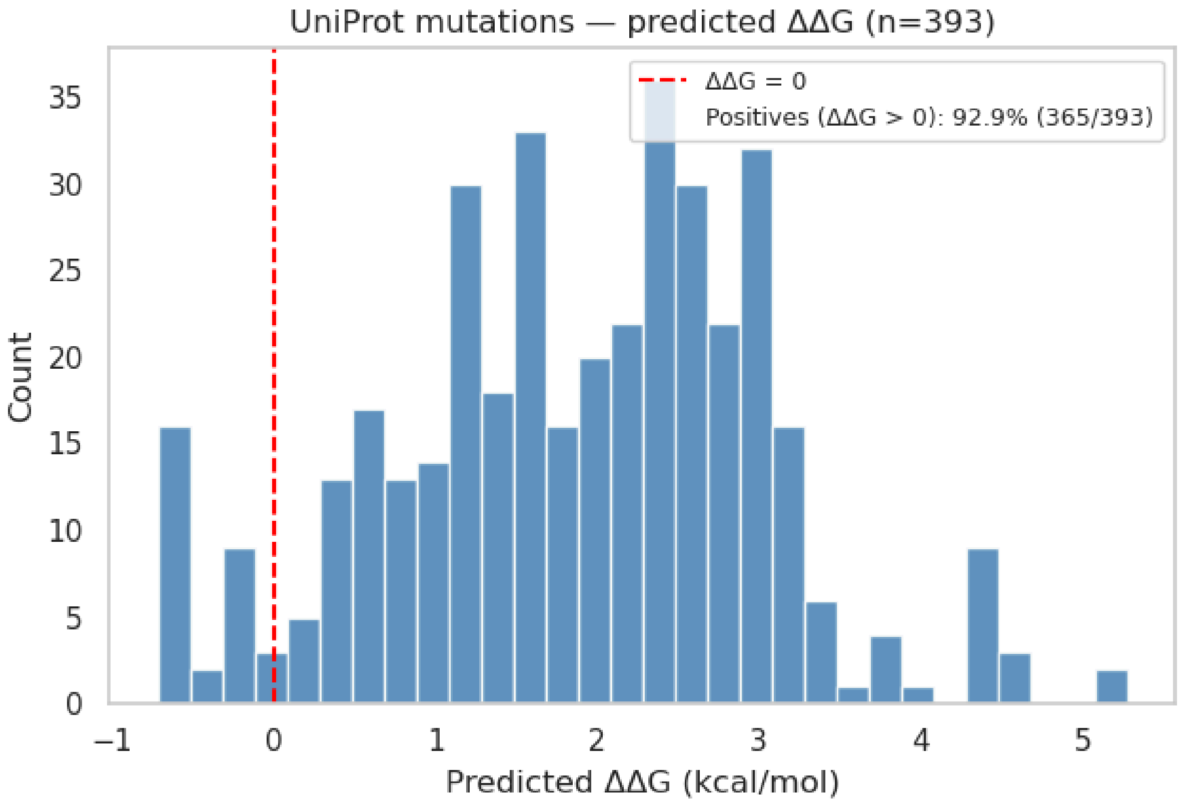


**Supplementary Figure S19. Distribution of predicted ΔΔG values for an independent set of RNA-binding-disrupting mutations.** Histogram of the ΔΔG values predicted by the Gradient Boosting regression model for the set of mutations collected from UniProt and described in the main text as experimentally validated to abolish or strongly impair RNA-binding activity. The histogram also reports the percentage of mutations predicted to have a positive ΔΔG. Notably, this percentage is ~93%, in agreement with the expected behavior, as these mutations are experimentally known to disrupt protein–RNA interactions and are therefore expected to destabilize binding. Importantly, these variants were not used during model training and include mutations from diverse proteins, as well as cases containing multiple amino acid substitutions, supporting the ability of the HERCULES-derived profiles to generalize beyond the benchmark dataset. Positive predicted ΔΔG values indicate weaker binding, consistent with the expected effect of mutations experimentally annotated to impair RNA binding.

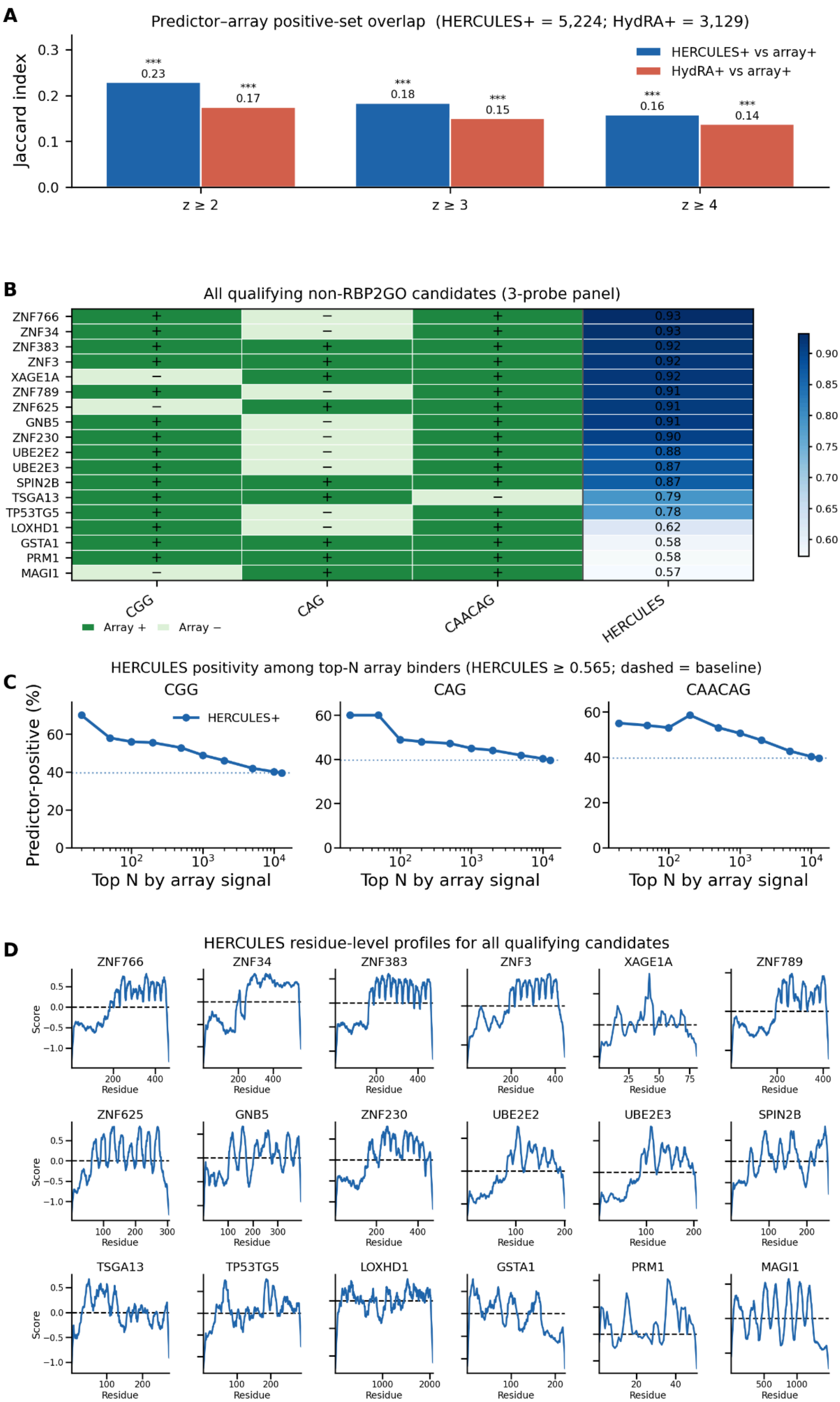
A
Predictor–array positive-set overlap (HERCULES+ = 5,224; HydRA+ = 3,129)
Jaccard index
HERCULES+ vs array+
HydRA+ vs array+
*** 0.23
*** 0.17
*** 0.18
*** 0.15
*** 0.16
*** 0.14
z ≥ 2
z ≥ 3
z ≥ 4
B
All qualifying non-RBP2GO candidates (3-probe panel)
ZNF766
ZNF34
ZNF383
ZNF3
XAGE1A
ZNF789
ZNF625
GNB5
ZNF230
UBE2E2
UBE2E3
SPIN2B
TSGA13
TP53TG5
LOXHD1
GSTA1
PRM1
MAGI1
CGG
CAG
CAACAG
HERCULES
Array +
Array −
HERCULES score
C
HERCULES positivity among top-N array binders (HERCULES ≥ 0.565; dashed = baseline)
CGG
CAG
CAACAG
Predictor-positive (%)
HERCULES+
Top N by array signal
D
HERCULES residue-level profiles for all qualifying candidates
ZNF766
ZNF34
ZNF383
ZNF3
XAGE1A
ZNF789
ZNF625
GNB5
ZNF230
UBE2E2
UBE2E3
SPIN2B
TSGA13
TP53TG5
LOXHD1
GSTA1
PRM1
MAGI1
Score
Residue

**Supplementary Figure S20. Protein-array validation of HERCULES-predicted non-catalogued RBP candidates. (A)** Jaccard overlap between HERCULES-positive proteins (global score > 0.565) or HydRA-positive proteins (HydRA score > 0.893) and array-positive proteins at increasing robust-z thresholds ($z \geq 2, 3, 4$; any of the CGG, CAG23, CAACAG probes). All overlaps significantly exceed random expectation (one-sided hypergeometric test; *** $p < 0.001$). **(B)** Heatmap of all 18 qualifying proteins absent from RBP2GO that passed the candidate-selection criteria: HERCULES global score > 0.565, array positivity on at least two of three repeat-RNA probes (CGG, CAG23, CAACAG; robust $z \geq 3$ per probe), and broad residue-level HERCULES signal (fraction of profile residues with score > 0 ≥ 0.5). Rows are ranked by descending HERCULES global score. Green cells denote array-positive (+) or array-negative (−) calls for each probe; the HERCULES column shows global scores. **(C)** Percentage of proteins predicted as positives by HERCULES, among top-N array binders, shown separately for CGG, CAG23 and CAACAG. Proteins on each probe were ranked by mean array signal; the percentage of HERCULES-positive proteins (score ≥ 0.565) within the top N binders is plotted against N (log scale). Dashed horizontal lines indicate the proteome-wide HERCULES-positive baseline on that probe. **(D)** HERCULES residue-level profiles for all 18 qualifying candidates. Each panel shows per-residue HERCULES scores along the sequence; the dashed line marks HERCULES z-score = 0. Profiles illustrate the extent of predicted RNA-binding signal across each candidate sequence. Protein microarrays were probed with synthetic RNAs enriched for CGG, CAG23 and CAACAG repeats. Array positivity was defined per probe using robust $z \geq 3$. Candidate selection and scoring criteria are detailed in **Supplementary Table S8** and **Methods**.
